# Choroid plexus sex differences in secretory signalling and immune compartments

**DOI:** 10.1101/2025.08.15.670566

**Authors:** Karol Kaiser, Violeta Silva-Vargas, Luca von Allmen, Thomas Sakoparnig, Fiona Doetsch

## Abstract

The choroid plexus (ChP)-cerebrospinal fluid (CSF) axis is emerging as a key regulator of the brain environment. The ChP is a multi-functional structure, which forms the blood-CSF barrier, and acts as a sensor of inputs from the periphery and the brain, dynamically responding by modulating its secretome into the CSF. However, sex differences in the ChP have been little explored. Molecular profiling of the adult mouse ChP at the transcriptomic and proteomic levels revealed sex differences across multiple cell types, with immune signatures enriched in females. Interestingly, border associated macrophages showed distinct sex differences in the stromal and epiplexus compartments. We further uncovered sex differences in ChP secretory signalling, with functional differences between males and females. Finally, the human LVChP showed largely similar sex differences. Together, our findings highlight that sex differences may play an important role in ChP function in both health and disease.

## Introduction

The choroid plexus (ChP) and cerebrospinal fluid (CSF) are emerging as key players in central nervous system homeostasis, brain physiology and disease. The CSF flows through the brain ventricular system, and serves both as a delivery route for diverse factors and removal of waste byproducts from the brain. The bulk of the CSF is produced by the ChP, a structure found in each brain ventricle, comprising a monolayer of epithelial cells surrounding a fenestrated vascular plexus and stromal core (Fig. 1a). ChP epithelial cells are polarized and inter-connected by tight junctions to form the blood-CSF-barrier, and provide a bi-directional transport system between the CSF and blood (Saunders et al 2023). The ChP also acts as an enzymatic barrier, and plays a key neuroprotective role by detoxification and active clearance of metabolites, as well as immunosurveillance by diverse immune cells, including border-associated macrophages (BAMs) (Ghersi-Egea et al 2018).

**Fig 1.**
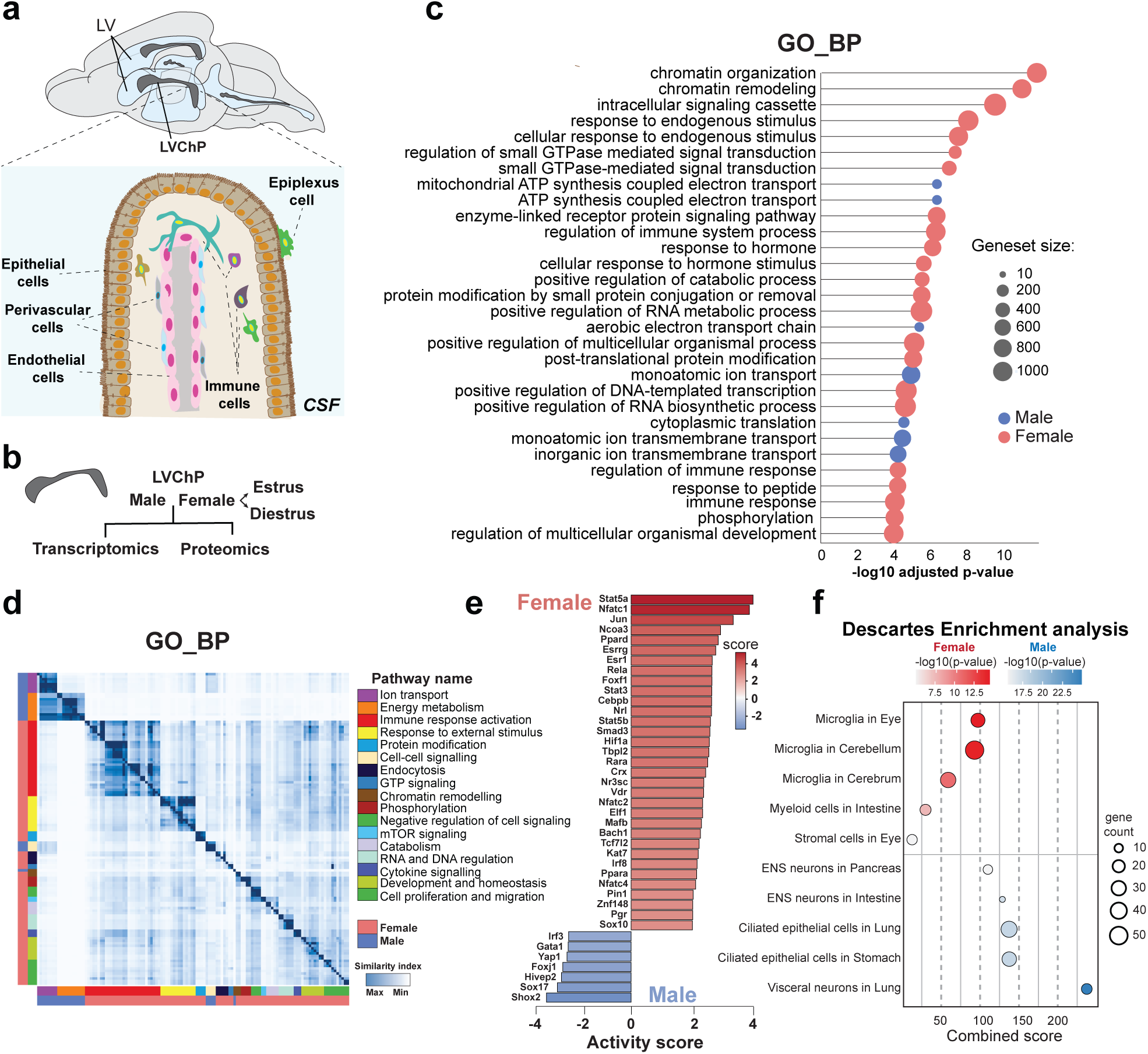
Global transcriptomic analysis of male and female lateral choroid plexus. **a** Schema of adult mouse brain showing location of lateral ventricles (LV) and lateral ventricle choroid plexus (LVChP). CSF flows through the brain ventricles. Magnification below shows schema of cell types in the LV ChP. Epithelial cells interconnected by tight junctions form the blood-CSF barrier, which surrounds the stromal compartment containing fenestrated blood vessels made of endothelial cells and perivascular cells, as well as distinct types of immune cell types. Epiplexus cells are located on apical side of epithelial cells. **b** Experimental design. Whole male and female (at estrus and diestrus stages of the estrous cycle) LVChPs were collected and sequenced for bulk transcriptomic analysis or for proteomics. **c** Lollipop plot showing top 30 statistically most significant pathway terms from Gene Ontology – Biological process database analysis for male and female LVChP transcriptome using the GOAT algorithm. **d** Gene set similarity heatmap with Gene Ontology – Biological process pathway terms grouped into categories. Color labels in legend refer to overarching group categories and their enrichment in males (blue) or females (red). Similarity index indicates pathway term similarity. **e** Inferred differential activity for top 40 most differentially activated transcription factors (TFs) in females and males. The x-axis represents the activity score, a measure of differential TF activity. Red bars highlight TFs with higher predicted activity in females, and blue bars in males. **f** Dot plot showing the top 5 enriched cell types from the Descartes_Cell_Types_and_Tissue_2021 database. Female-upregulated pathways (red) and male-upregulated pathways (blue) were identified based on positive and negative *z*-scores, respectively, calculated by the GOAT algorithm and ranked by raw *p*-values (*-log₁₀(p)* shown as color scale).

Importantly, the ChP is a source of a large repertoire of biologically active molecules in the CSF. The ChP secretome contains factors actively produced by ChP cells as well as some derived from the blood. In addition to its complexity, the ChP secretome is highly dynamic, thereby contributing to changes in CSF composition (Ghersi-Egea et al 2018). In the adult, the ChP secretome changes over short time scales, including diurnal changes and in response to acute insults including inflammation, as well as throughout the lifespan (Saunders et al 2023). As such, the ChP is garnering increasing attention due to its emerging role as a hub that integrates signals from the CSF and bloodstream to modulate its secretome in response to various physiological signals. ChP-derived factors can be delivered to both local and distant brain areas via the CSF to regulate multiple cell types and processes, such as adult neural stem cell regulation, circadian clock regulation and behaviour, including olfaction and memory (Baruch et al 2014, Liu et al 2023, Myung et al 2019, Phillipps et al 2025, Silva-Vargas et al 2016, Taranov et al 2024), as well as neural plasticity during critical periods (Spatazza et al 2013). In addition, several pathological conditions, including hydrocephalus, Alzheimer’s disease (AD), Multiple Sclerosis (MS), schizophrenia, Parkinson’s disease, Amyotrophic lateral sclerosis (ALS), Autism spectrum disorder (ASD) and Psychosis have been associated with ChP impairment (Bothwell et al 2019, Brunner et al 2025, Choi et al 2022, Courtney et al 2025, Husag et al 1984, Jeong et al 2023, Levman et al 2018, Lizano et al 2019, Rodriguez-Lorenzo et al 2020). Therefore, understanding the molecular mechanisms that control ChP functions is critical to understanding its potential role in homeostasis and disease.

Despite its central role in brain physiology, little is known about sex differences in the ChP. Sex differences can play key roles in disease and different physiological processes. Interestingly, sex hormone receptors are expressed by the ChP (Santos et al 2017). Previous studies showed sex differences in the ChP transcriptome and proteome in gonadectomised rats (Quintela et al 2013, Quintela et al 2016) and that CSF production differs between sex in mice (Liu et al 2020).

Here we examined the adult mouse lateral ventricle ChP (LVChP) in males and females and uncover important sex differences at the level of the transcriptome, proteome, and secretome, with females having an enriched immune signature. Secretory signalling exhibits functional differences between males and females. Moreover, BAMs show important sex differences across compartments, with stromal macrophages more abundant in the female LVChP, and epiplexus cells in males. Strikingly, similar sex differences are found in the human ChP. Together, our findings highlight the importance of sex differences in understanding the biology of the ChP in health and disease.

## Results

### Molecular profiling of the lateral ventricle choroid plexus reveals sex differences

To gain insight into sex differences in the choroid plexus, we performed transcriptomic and proteomic profiling of the LVChP of male and female adult mice (Fig. 1b). To control for hormonal fluctuations, we collected tissue during two distinct estrous cycle phases; estrus and diestrus, which exhibit very different hormone profiles (Ajayi & Akhigbe 2020).

We first compared the transcriptomes of males and females independent of estrous cycle state to assess overall differences between sexes. Interestingly, global gene set enrichment analysis showed a higher number of enriched pathways from the GO Biological process database (24 of the top 30 most significant pathway terms) upregulated in females as compared to males (Fig. 1c) (Supplementary Table 1). In females, about one third of all biological processes were related to immune response, with the others corresponding to response to stimuli, RNA/DNA regulation, posttranslational modification and degradation, phosphorylation, vesicle transport, and endocytosis (Fig. 1d). In contrast, males showed upregulation for processes related to ion transport, ion channels, energy metabolism, cell-cell signalling processes and translation (Fig. 1c, d). Molecular function and cellular compartment filtering highlighted similar corresponding categories (Supplementary Fig. 1a-d).

To evaluate whether sex-differences reflect differences in underlying gene regulatory networks, we inferred transcription factor activity from male and female LVChP transcriptomes. Females exhibited a higher number of differentially active TFs among the top 40 identified, as compared to males (Fig. 1e). TF activity for immune regulation and signalling was high in females, consistent with the large number of immune related GO biological pathways in females. For Stat5a, an immune regulator (Yao et al., 2006), targets annotated as activated were predominantly upregulated in females, whereas targets annotated as repressed were predominantly upregulated in males (Supplementary Fig. 1e). Interestingly, nuclear receptor TF activity, including sex hormone receptors, was also enriched in females (Fig. 1e). In males, cilia and mechanotransduction-related TFs were among those with the highest TF activity, as well as some immune-related TFs (Fig. 1e). Notably, cell identity inference with Enrichr revealed microglia as the most represented cell type in females, whereas epithelial and neuronal signatures were enriched in males (Fig. 1f).

Together, this global analysis highlights that the LVChP transcriptome exhibits sex differences, with immune-related pathways emerging as potential key players in females.

### Sex differences occur across multiple choroid plexus cell types

We next performed differential gene expression analysis between males and females, which revealed 154 differentially expressed genes (DEGs, adjusted p-value < 0.05) between sexes. As expected, samples segregated by sex (Supplementary Fig. 2a), and sex chromosome genes were among the top DEGs (Supplementary Fig. 2b, c, Supplementary Table 2), including Kdm6a, Ddx3x which escape X inactivation (Berletch et al 2015). DEG analysis revealed 120 autosomal genes, 55 of which were upregulated in males, while 65 genes were upregulated in females (Fig. 2a; Supplementary Table 2), which we focused on for analysis.

**Fig 2.**
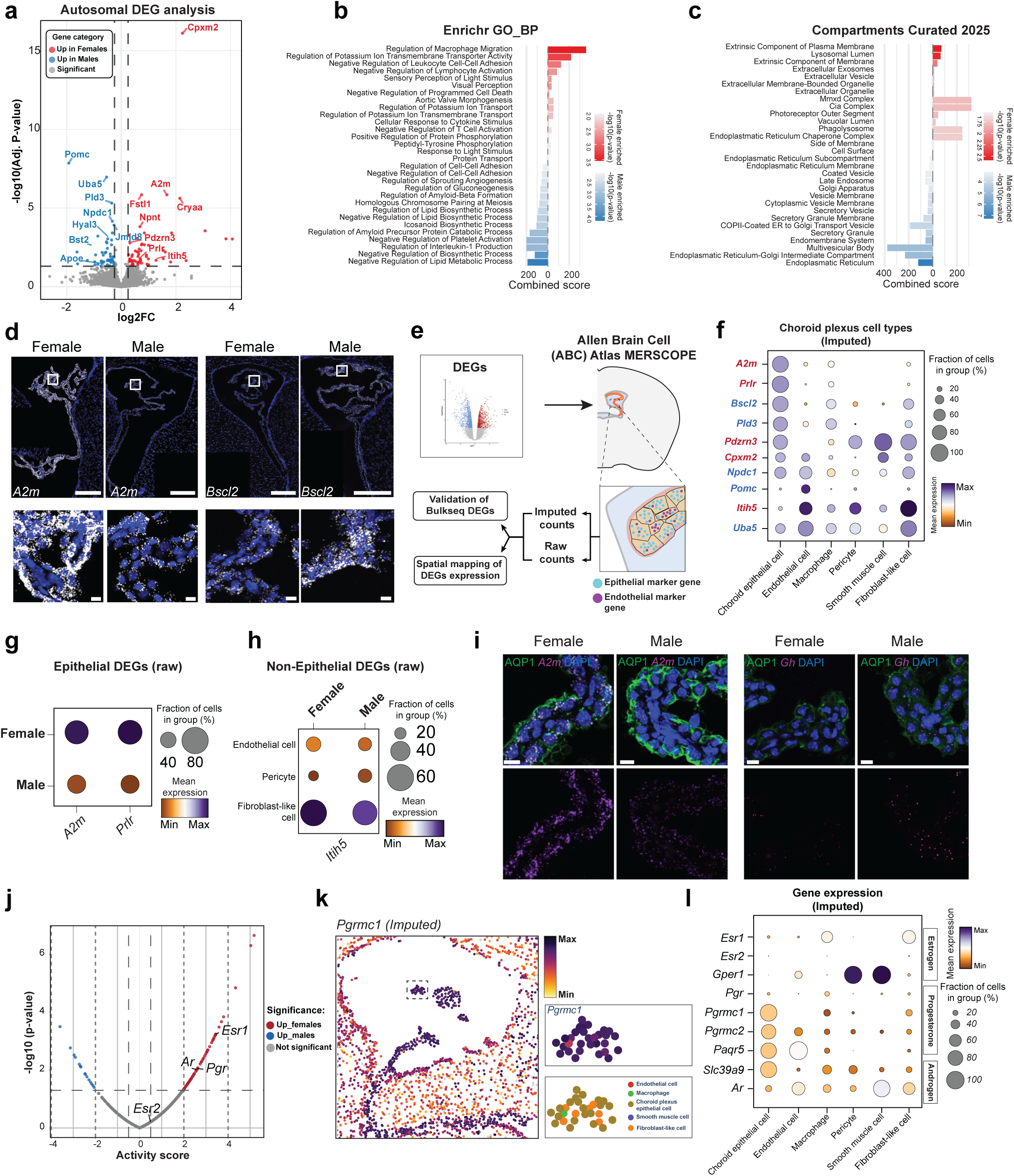
Sex differences in the LVChP transcriptome. **a** Volcano plot of differentially expressed genes (DEGs) in the male and female LVChP. Significance thresholds: adjusted p-value (padj) < 0.05, ∣log2FC∣>0.25. The top 10 most significant genes are highlighted for females (red) and males (blue), with some additional manually added DEGs displayed. **b, c** Pathway enrichment analysis for male and female DEGs using Gene Ontology (Biological Process - BP (**b**) & Cellular compartment – CC (**c**)) databases. Split bar plots show the top pathways, selected based on p-value and then ranked by combined score (x-axis). Red and blue gradients indicate enrichment (–log10(p-value)) in females and males, respectively. **d** *In situ* hybridization (RNAscope) for *A2m* (female upregulated) and *Bscl2* (male upregulated) DEGs. Insets show higher-magnification images. Scale bar: 250µm. Inset scale bar: 10 µm. **e** Workflow for spatial transcriptomic analysis of DEGs in female and male LVChP using publicly available MERSCOPE dataset from Allen Brain Cell Atlas. **f** Dot plot of imputed cell type expression of top DEGs in LVChP. Red font highlights female-up-regulated, blue font male-enriched DEGs. **g** Dot plot of raw expression differences from MERSCOPE data for epithelial-enriched DEGs. **h** Dot plot of raw expression differences for *Itih5,* a non-epithelial enriched DEG. **i** Immunostaining for AQP1, an apical ChP epithelial marker, combined with *in-situ* hybridization for *A2m* or *Gh*. Scale bar: 10µm. **j** Volcano plot showing sex-hormone receptor related transcription factor activity. Activity scores represent differential activity between females and males (x-axis). Statistical significance is shown as −log10(p-value) (y-axis). Significance thresholds used: p-value < 0.05 and absolute activity score > 0.5. Red dots: significantly up-regulated in females; blue dots: significantly up-regulated in males; grey dots: not significant. **k** Spatial UMAP plot showing imputed spatial expression for *Pgrmc1* in LVChP versus adjacent regions. Insets show highest expression in LVChP epithelial cells. **l** Dot plot of imputed cell type expression of female and male sex hormone-related receptors in LVChP.

The top 15 GO Biological pathways for female DEGs were related to regulation of macrophage migration, sensory perception of light stimulus, immune cell adhesion and activation, potassium ion transport, and cellular response to cytokine stimulus. Top pathways enriched in males were related to lipid metabolism, Interleukin production, platelet activation, amyloid processing and cell-cell adhesion (Fig. 2b). Cell compartment analysis highlighted terms related to secretion, but differing between sex, including secretory granule, multi-vesicular body and secretory vesicles in males, and extrinsic component of membrane, extracellular vesicles, and phagolysosome in females (Fig. 2c). Cell type inference again highlighted macrophages as the most enriched cell type in females, and epithelial-related cells in males (Supplementary Fig. 2d).

We confirmed the differential expression of some male-enriched (*Bst2, Hyal3, Bscl2, Gh*) and female-enriched (*A2m, Itih5, Cpxm2*) genes using qPCR (Supplementary Fig. 2e) or RNAscope (Fig. 2d, i, Supplementary Fig. 2f). Interestingly, some DEGs (*A2m, Cpxm2, Bscl2*) were highly enriched in the ChP relative to the rest of the brain (Fig. 2d, Supplementary Fig. 2f). Importantly, RNAscope revealed that DEGs were expressed in different spatial patterns and in distinct ChP cell types, in both the epithelial layer and in the stromal compartment, which includes endothelial cells, pericytes, macrophages, smooth muscle cells, and fibroblast-like cells (Fig. 2d, Supplementary Fig. 2f). To gain further insight into which ChP cell types express DEGs, we used the spatial MERSCOPE transcriptomic data from the Allen Brain Cell Atlas (Fig. 2e). This spatial analysis likewise revealed different expression patterns across ChP cell types for DEGs, with some predominantly enriched in epithelial cells (*A2m, Prlr, Bscl2, Pld3*), others in stromal cell types (*Itih5, Pomc, Uba5*) and some in both (*Cpxm2, Pdzm3, Npdc1*) (Fig. 2f). Analysis of the raw MERSCOPE data for available probes by sex confirmed the higher expression in females of *A2m* and *Prlr* in epithelial cells and *Itih5* in stromal cells in females (Fig. 2g, h). The epithelial enrichment of *A2m* and *Gh* was also validated by immunostaining for the apical epithelial marker Aquaporin 1 (Aqp1) (Fig. 2i). Finally, female upregulation of *A2m* expression was also detected in the third and fourth ventricle choroid plexuses (Supplementary Fig. 2g, h).

Our TF activity mapping showed that nuclear sex hormone receptors were among the top predicted differentially active TFs in the female choroid plexus (Fig. 1e, 2j). Indeed, the choroid plexus expresses receptors for sex hormones including estrogen, progesterone and androgens (reviewed in Santos 2017). Analysis of female spatial MERSCOPE data from the Allen Brain Cell Atlas confirmed sex hormone receptor expression in the ChP, and strikingly, revealed that several were highly enriched in the ChP compared to other brain areas (*Paqr5, Esr1, Pgrmc1)* (Fig. 2k, Supplementary Fig. 3a, b). Moreover, within the ChP itself, sex hormone receptor expression was not uniform, rather showed distinct profiles in different ChP cell types (Fig. 2k, l, Supplementary Fig. 3b). Some receptors for the same sex hormone were more broadly expressed across ChP cell types (*Pgrmc 2, Slc39a9, Ar*), whereas others were more selectively localized, with *Esr1* enriched in macrophages and fibroblast-like cells, and *Gper1* in endothelial cells, pericytes and smooth muscle cells, (Fig. 2k, l; Supplementary Fig. 3b). Interestingly, membrane-associated progesterone and testosterone receptors were more enriched in epithelial cells, whereas sex hormone nuclear receptors were more enriched in stromal compartment cells (Fig. 2l).

The diverse expression patterns of sex hormone receptors suggests that the choroid plexus may be sensitive to fluctuations in sex hormones levels. We therefore analyzed how the LVChP transcriptome changes with the estrous cycle, directly comparing estrus and diestrus females. DEG analysis revealed more genes upregulated in estrus (Supplementary Fig. 3c). Gene ontology (GO) pathways upregulated in estrus included categories related to ion channels, synaptic transmission, and calcium signalling (Supplementary Fig. 3d-f). In diestrus, pathways related to blood vessels and extracellular matrix (ECM) were enriched (Supplementary Fig. 3d-f). Moreover, TF activity analysis suggests that different gene regulatory networks are active in the LVChP in estrus and diestrus (Supplementary Fig. 3g).

Together these findings highlight that DEGs are distributed across multiple ChP cell types in males and females, and that the ChP may be especially attuned to changes in hormone levels over the estrous cycle, with distinct ChP cell types responding to different hormones.

### Sex differences in choroid plexus secretory signalling

The ChP is a key source of factors that can be secreted into the CSF, thereby changing CSF composition and modulating brain physiology. Given the pronounced differences between males and females in our LVChP transcriptome analysis, with many terms associated with secretion upregulated in male and females, we explored sex differences in the ChP secretome. We first analyzed sex differences in the LVChP proteome (Fig. 3a). Mass spectrometry of the whole ChP identified 777 differentially expressed peptides (DEPs, q-value < 0.05). More differentially enriched proteins were detected in females as compared to males (709 versus 68 proteins) (Fig. 3b; Supplementary Table 3). Global pathway enrichment analysis of the top 25 differentially expressed pathways of the proteome uncovered a greater number upregulated in females. These included pathways related to fluid secretion, vesicular and cellular transport, energy building metabolism, copper ion homeostasis, protein translation and protein folding (Fig. 3c). In males, top enriched pathways were associated with coagulation, fibrinolysis, vesicle fusion and signalling cascades (Fig. 3c). Notably, some DEGs, including *A2m, Cpxm2, Prlr, Dhx32, Niban1, Pepd, Cryz,* in females and *Ptgds* in males, were also differentially enriched at the proteome level (Fig. 3b), whereas others were not, suggesting potential post-transcriptional regulation in the ChP.

**Fig 3.**
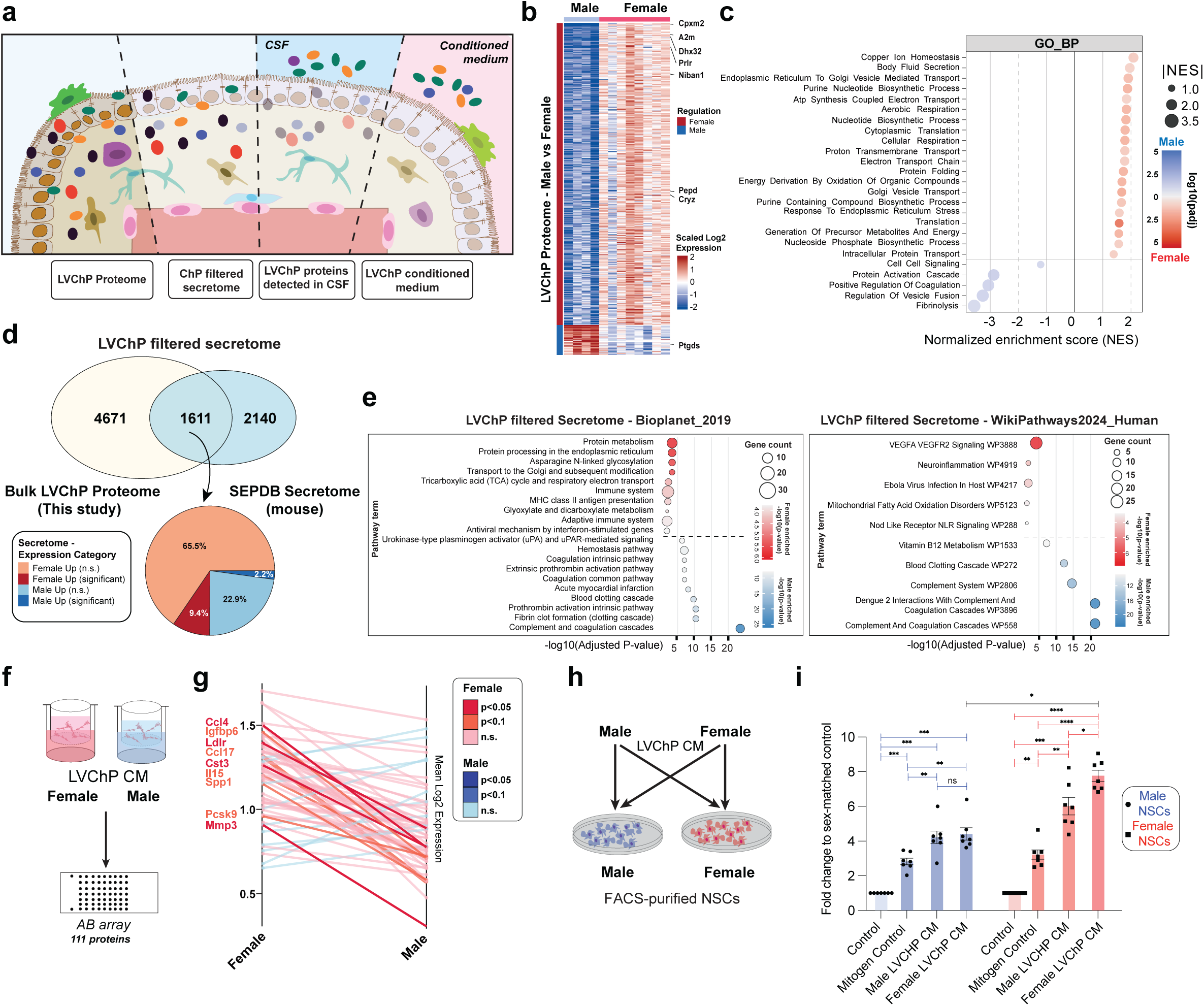
Functional analysis of sex differences in the LVChP secretome. **a** Schema showing different analyses of the LVChP proteome and secretome. First, female and male LVChP proteomes were defined using mass spectrometry (left), followed by filtering for putative secreted proteins (from SEPDB database) (second from left), and for proteins detected in mouse CSF (second from right). Finally LVChP conditioned medium was analyzed using antibody arrays. **b** Heatmap showing differentially enriched proteins in male and female LVChP. Highlighted proteins were also enriched as DEGs in transcriptomic analysis. Significance thresholds adjusted p-value (padj) < 0.05, ∣log2FC∣>0.25. **c** Split dot plot showing top 25 statistically most significant pathway terms from Gene Ontology Biological Process - BP database analysis for male and female LV ChP proteome using fGSEA. **d** Venn diagram showing overlap between total LVChP proteome (male and female LVChP combined) and SEPDB (published curated database of secreted proteins). Pie chart showing proportions of overlapping putative secreted proteins enriched in male and female LVChP proteome. **e** Combined dot plots showing pathway enrichment analysis for DEPs using Bioplanet_2019 or WikiPathways2024_human databases. Red and blue gradients (–log10(p-value)) indicate enrichment in females and males, respectively. **f** Schematic representation of LVChP conditioned medium (CM) experimental design. CM from male or female LVChP explants were analyzed by antibody arrays. **g** Slope plot showing enrichment of LVChP secreted proteins detected in antibody arrays. Red lines highlight female enriched proteins, blue lines male enriched proteins. Darker color shade indicates statistically significant enriched proteins (p value < 0.05 or 0.1). **h** Schematic showing NSC experiment. **i** Quantification of NSC activation with male and female LVChP conditioned medium. Statistics: 2-way ANOVA and Šídák’s multiple comparisons test.

To examine the LVChP secretome, we applied three complementary approaches (Fig. 3a). First, to identify LVChP secreted proteins, we filtered the bulk LVChP proteome against an annotated secreted proteins database (SEPDB), which revealed 1611 LVChP potential secreted proteins (Fig. 3a, d and Supplementary Table 3). Females again had more differentially enriched secreted proteins (9.5%) than males (2.2%) (Fig. 3d). Interestingly, many female enriched proteins were related to MHC antigen presentation, adaptive immunity, anti-viral response, and protein folding and modification, and neuroinflammation (Fig. 3e). In contrast, male enriched secreted proteins were primarily associated with complement and coagulation pathways (Fig. 3e). Second, we compared our LVChP proteome with that of a publicly available mouse CSF proteome dataset (Fig. 3a, Supplementary Fig. 4a). The majority of proteins detected in the CSF proteome (87%) overlapped with proteins in the LVChP (390 proteins) (Supplementary Fig. 4a), with a 12.6% and 6.2% enrichment in female and male CSF respectively, confirming the ChP as an important contributor to the CSF. Combined GO pathway enrichment analysis revealed that female enriched CSF proteins were associated with neuroinflammation, antigen presentation and metabolism, whereas male enriched proteins were related to the complement and coagulation pathways (Supplementary Fig. 4b).

Finally, we investigated sex differences in the LVChP secretome by preparing conditioned medium from male and female LVChP and profiling 111 proteins using antibody arrays (Fig. 3a, 3f). This identified 63 detected proteins (Fig. 3g, Supplementary Table 3) many of which were enriched in females (Fig. 3g, Supplementary Fig. 4c). These included signalling and matrisome proteins, cytokines, metabolic regulators and protease inhibitors (Supplementary Fig. 4c, Supplementary Table 3). The LVChP secretome also exhibited changes in estrus and diestrus (Supplementary Fig. 4d, e).

ChP factors are secreted into the CSF. The ventricular-subventricular zone (V-SVZ), the largest stem cell niche in the adult brain, lies adjacent to the lateral ventricles and is directly exposed to CSF factors (Delgado et al 2014, Delgado et al 2021, Lepko et al 2019, Sawamoto et al 2006, Silva-Vargas et al 2016) (Supplementary Fig. 4f). Factors in the CSF can affect many cell types in the V-SVZ, including multi-cilated ependymal cells that line the brain ventricles and neural stem cells that directly contact the CSF, as well lineage cells and other niche cells in the underlying tissue that do not directly contact the CSF (Delgado et al 2014, Pastrana et al 2009, Sawamoto et al 2006). Ligand receptor pairing analysis revealed that the ChP provides a rich repertoire of secreted ligands that can interact with many V-SVZ cell types based on the expression profile of corresponding receptors (Supplementary Fig. 4g).

We previously discovered that adult neural stem cells (NSCs) are particularly sensitive to changes in the ChP secretome with aging (Silva-Vargas et al 2016). To directly assess functional differences between male and female ChP-derived factors, we assayed clone formation by FACS-purified male and female NSCs cultured with either sex matched or sex-swapped LVChP conditioned medium (LVChP secretome (LVChPsec) (Fig. 3h, i). As we previously reported (Silva-Vargas et al 2016), LVChPsec promoted greater clone formation than standard mitogens (Fig. 3i). Strikingly, LVChPsec showed sex differences. Female NSCs formed more clones in response to female LVChP secretome than to male LVChP secretome (Fig. 3i). Moreover, female LVChP-derived factors were more potent on female NSCs than on male NSCs (Fig. 3i). Notably, there was no significant difference in baseline activation of male and female NSCs when cultured with EGF alone (Fig. 3i).

Together, these results reveal pronounced sex differences in ChP secretory signaling that influence adult NSCs and may have broader implications for cell populations throughout the brain.

### Border associated macrophages show compartment-specific sex differences

One important role of the ChP is immune surveillance. In our transcriptomic and proteomic analyses, immune-related signatures were highly enriched in females. Microglia/macrophages were the major over-represented cell type in females (Fig. 1f, Supplementary Fig. 2d), prompting us to investigate sex differences within this cell population. Choroid plexus macrophages belong to a unique class of cells called BAMs, which are distinct from microglia in the brain parenchyma.

BAMs can be visualized by immunostaining for IBA1 (ionised calcium-binding adapter molecule), CX3CR1 or CSFR1, which are largely overlapping (Supplementary Fig. S5 a-b). We first quantified total BAMs in whole mount preparations of the LVChP. BAMs were more abundant in females in IBA1-immunostained whole mount preparations (Fig. 4a, b), and in CX3CR1-GFP mice (Supplementary Fig. 5c). Distinct BAM morphologies were observed, including amoeboid, bipolar, intermediate cells and stellate cells (Choi et al 2021, Karperien et al 2013) (Fig. 4c). Significantly higher numbers of intermediate and stellate BAMs were present in the female LVChP as compared to males, with no difference in the density of amoeboid or bipolar BAMs (Fig. 4c). Many stellate stromal BAMs extended podocyte-like processes that contacted the vasculature (Supplementary Fig. 5d). Quantification of Iba1 immunostaining in coronal sections confirmed the increased total number of BAMs in females (Supplementary Fig. 5e).

**Fig. 4.**
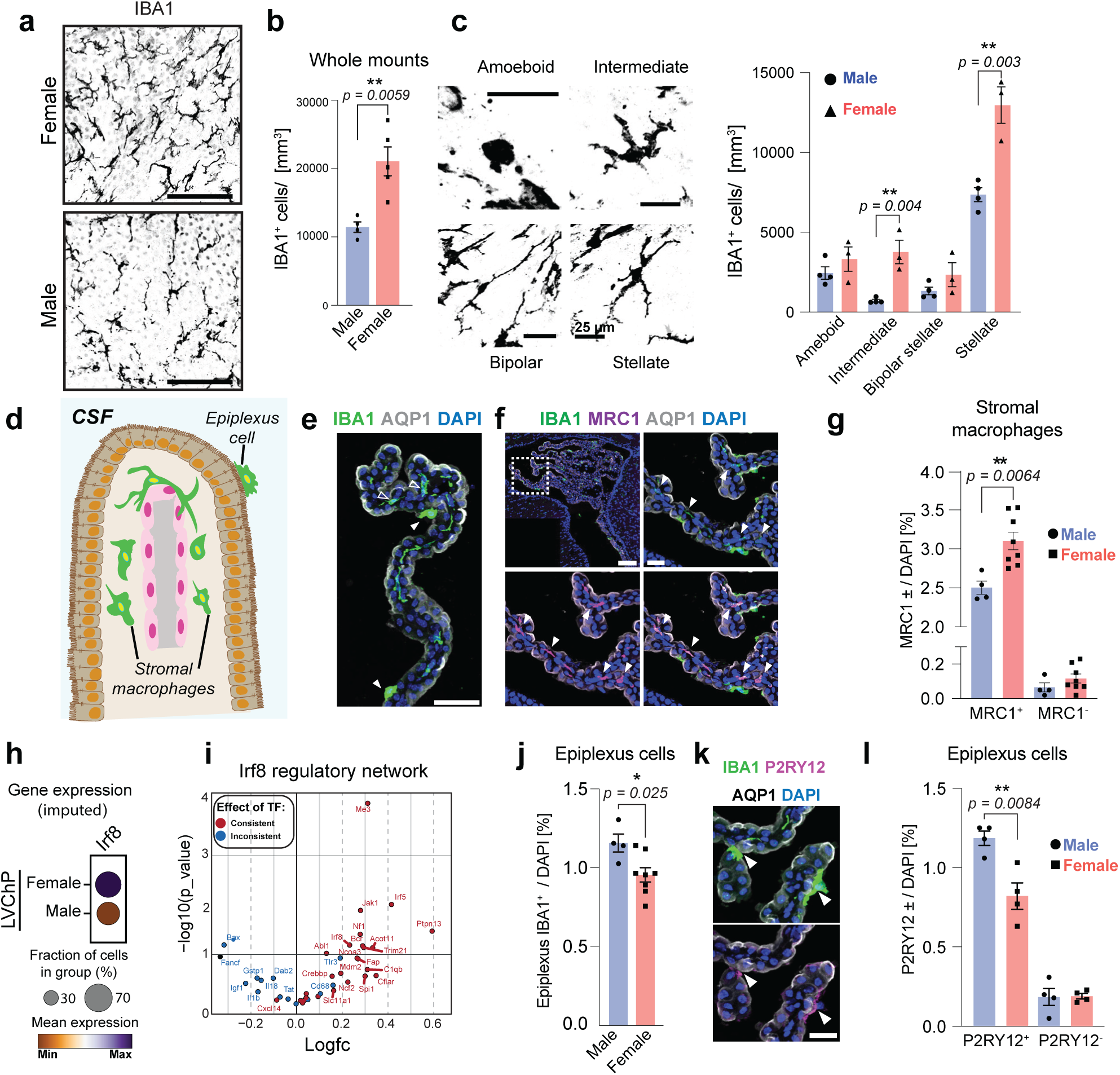
Sex differences in border associated macrophage compartments. **a** Representative image of whole mount preparation immunostained for IBA1 (black) in female (top) and male (bottom) LVChP. **b** Quantification of total IBA1+ BAMs in male and female whole mounts. **c** Representative images of different IBA1+ macrophage morphologies and their quantification. **d** Schema of the LVChP BAM compartments (stromal and epiplexus macrophages). **e** Representative image of a cross-section immunostained for IBA1 (green), AQP1 (white) and DAPI (blue) showing epiplexus macrophages (white arrowheads) and stromal macrophages (empty arrowheads). Scale bar: 50µm. **f** Representative image of a cross-section immunostained for IBA1 (green), MRC1 (magenta), AQP1 (white) and DAPI (blue) showing stromal macrophages (white arrowheads). Box shows region shown at higher-magnification. Scale bar top left: 100µm. Other scale bars: 25µm. **g** Quantification of MRC1^+^ and MRC1^−^ stromal macrophages in male and female LVChPs. **h** Dot plot of sex expression differences of *Irf8* from imputed spatial transcriptomics data. **i** Volcano plot illustrating the differential expression of target genes for transcription factor Irf8 between female and male LV ChP. The x-axis represents the log-fold change (logFC), and the y-axis shows significance (−log10(p−value)). Genes are colored based on consistency between the gene’s regulation direction with the expected transcription factor’s mode of regulation: red for consistent, blue for inconsistent. **j** Quantification of total IBA1^+^ epiplexus macrophages in coronal sections. **k** Representative image of a cross-section immunostained for IBA1 (green), P2RY12 (magenta), AQP1 (white) and DAPI (blue) showing P2RY12^+^ epiplexus macrophages (white arrowheads). Scale bar: 25µm. **l** Quantification of P2RY12^+^ and P2RY12^−^ epiplexus cells in LVChP.

Choroid plexus BAMs reside in two distinct compartments, within the stroma (stromal macrophages), or on top of the apical surface of epithelial cells in direct contact with the CSF (epiplexus cells, also known as Kolmer cells) (Schwarze 1975, Van Hove et al 2019) (Fig. 4d). To examine sex differences in these compartments, we analyzed coronal sections of the LVChP immunostained with AQP1, which labels the apical surface of epithelial cells, allowing easy identification of epiplexus cells versus stromal macrophages (Fig. 4e). Stromal macrophages express mannose receptor C-type 1 (MRC1) (Van Hove et al 2019). MRC1+ immunostaining revealed that the vast majority of stromal IBA1+ cells express MRC1 (Supplementary Fig. 5f), and that MRC1+ stromal macrophages were more abundant in females (Fig. 4f, g). Spatial trancriptomics from the Allen Brain Cell Atlas of male and female adult mouse ChP further showed that the macrophage master regulator *Irf8* (Van Hove et al 2019) is highly expressed in BAMs and that Irf8 expression and its target genes were significantly elevated in females (Fig. 4h, i, Supplementary Fig. 5g). Unexpectedly, in the epiplexus compartment, a significantly higher number of epiplexus cells was found in male LVChP (Fig. 4j). Recent single-cell LVChP BAM profiling identified genes enriched in epiplexus cells, including *P2ry12* (Jordao et al 2019, Utz et al 2020, Van Hove et al 2019). Immunostaining for P2RY12 revealed that the majority (84%) of epiplexus cells were P2RY12+ IBA1+, and that these were more abundant in males (Fig. 5k, l; Supplementary Fig. 5h).

**Fig. 5.**
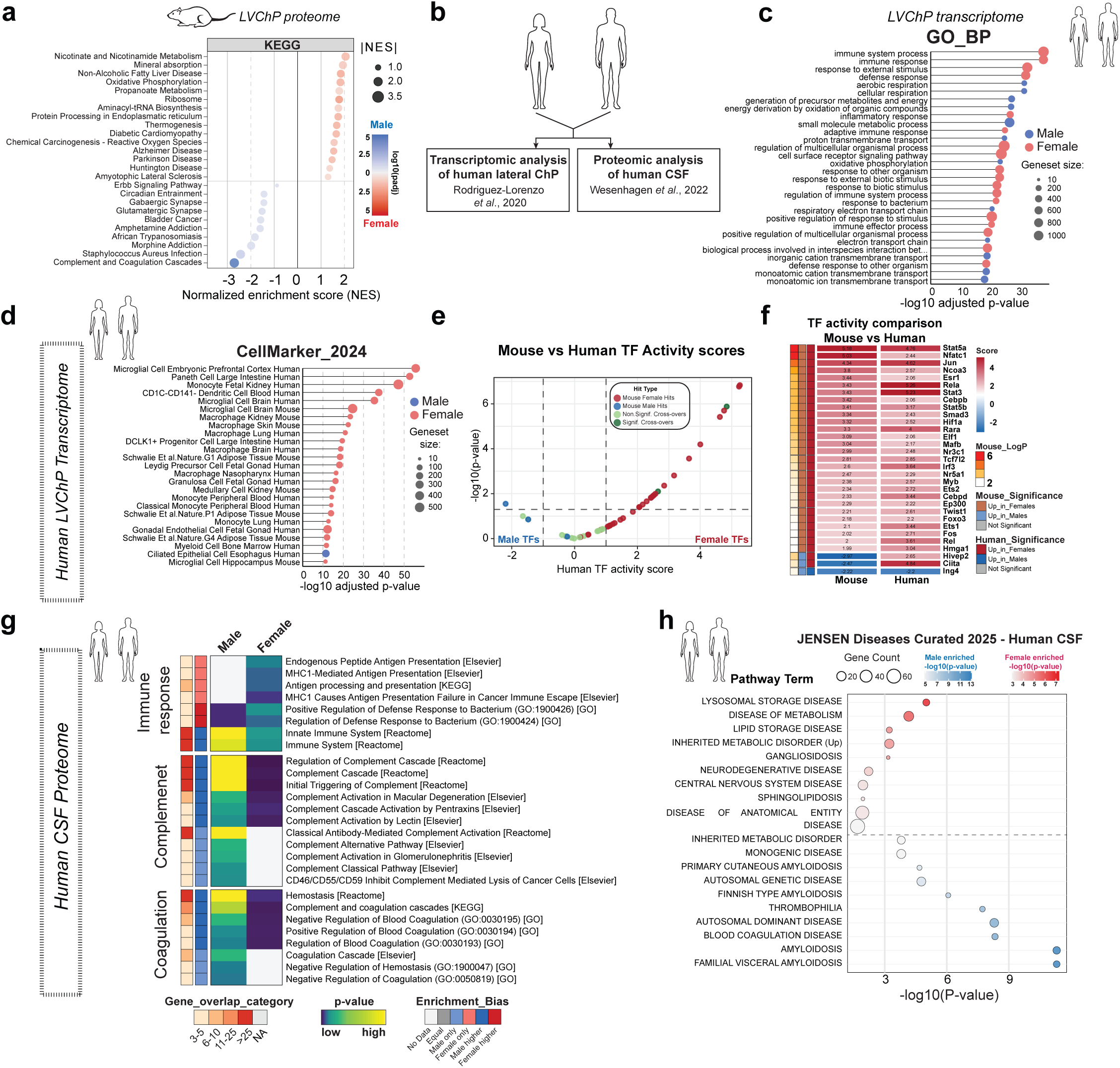
Functional similarity of mouse and human LVChP. **a** Split dot plot showing top 25 statistically most significant pathway terms from KEGG database analysis for mouse male and female LV ChP proteome using fGSEA. **b** Overview of human datasets used for comparative analysis with mouse LVChP transcriptome and proteome. **c** Lollipop plot showing top 30 statistically most significant pathway terms from GO:BP database analysis for human male and female LV ChP transcriptome using the GOAT algorithm. **d** Dot plot displays top 25 enriched cell types from CellMarker2024 database. Female-upregulated pathways (red) and male-upregulated pathways (blue) were identified based on positive and negative z-scores, respectively, calculated by the GOAT algorithm and ranked by raw p-values (-log₁₀(p) shown as color scale). **e** Volcano plot showing the comparison of top enriched TF hits from mouse bulkseq transcriptome compared to human female vs male LVChP transcriptomic analysis. Red dots are female TFs with enriched activity in both female mouse and human LVChP, blue dots in male mouse and human LVChP and green dots represent hits with opposite activity score between mouse and human LVChP transcriptome. **f** Heatmap showing male and female top TF hits and their activity score in both human and mouse LVChP. Thresholds used: p < 0.05, activity |score| > 1. **g** Heatmap showing enrichment between female and male CSF in terms related to the immune response, complement or coagulation pathway from Enrichr analysis of sex-enriched proteins in human CSF across various databases. **h** Combined dot plot showing top hits for enriched disease pathway terms from JENSEN_Diseases_Curated_2025 database. Dots with red and blue gradients (–log10(p-value)) indicate enrichment in female and male CSF proteome, respectively.

These findings reveal distinct BAM subpopulations in the LVChP with clear sex-dependent differences, with stromal macrophages predominating in females, and P2RY12+ epiplexus cells in males.

### Sex differences in mouse show parallels in human choroid plexus

The ChP is emerging as a key player in neurodevelopmental and neurodegenerative disease. To investigate whether ChP sex-enriched proteins are associated with disease, we performed pathway enrichment analysis of the mouse ChP proteome and secretome. This highlighted metabolic diseases and neurodegenerative diseases, including Parkinson’s, Alzheimer’s, Huntington’s and ALS in females, whereas complement and coagulation, infection and neurotransmitter related pathways were top enriched pathways in males (Fig. 3e, 5a, Supplementary Fig. 6a, b).

To assess whether the human ChP also exhibits sex differences, we analyzed publicly available human LVChP transcriptome (Rodriguez-Lorenzo et al 2020) and CSF datasets (Wesenhagen et al 2022) (Fig. 5b). Strikingly, global gene set enrichment analysis of human LVChP transcriptome revealed many similar patterns between mouse and human, with an enrichment for immune-related processes, extracellular matrix and response to external stimuli in females, and energy metabolism, respiration, amino acid biosynthesis, cilia and ion transport in males (Fig. 5c, Supplementary Fig. 6c-e). Macrophage and microglia signatures were the most enriched cell type category enriched in females, and epithelial cells in males in humans (Fig. 5d). Mapping of TF activity in human choroid plexus transcriptome also showed that of the top 40 transcription factors, the majority were active in females (Supplementary Fig. 6f). Strikingly, most of the mouse TF hits were also enriched in human ChP (Fig. 5e, f), again with several immune-related TFs in the female. Notably, for both mouse and human LVChP, the array of active TFs highlights the diversity of signals and changes in the peripheral and brain environments that the ChP is attuned to, including hormones, oxygen levels, immune status, and global metabolic inputs (Fig. 5f). Interestingly, disease analysis of human LVChP transcriptome showed different disease associations between sex (Supplementary Fig. 6g).

Comparison of male and female human CSF proteome showed key sex differences as well as a higher number of differentially expressed proteins in females (Supplementary Fig. 6h). Strikingly, as in mouse, comparison of human male and female CSF showed an enrichment of proteins in the immune response pathways in females and complement and coagulation pathways in males (Fig. 5g). Moreover, disease pathway analysis of human sex-enriched proteins in the CSF revealed significant hits in multiple neurodegenerative diseases, as well as lysosome storage diseases and lipid-related diseases in females (Fig. 5h). In males, upregulated pathways included thrombophilia, amyloidosis, thyroid and endocrine cancers, tauopathy and Alzheimer’s disease (Fig. 5h). Notably, differentially expressed proteins detected in the mouse LVChP secretome were also present in the human CSF and displayed the same direction of enrichment in males and females, including A2M, Cd9 and Pgk1 (female enriched) and complement associated proteins (F2, FGB and VTN (male enriched)) (Supplementary Fig. 6i-k).

In sum, the human choroid plexus also exhibits sex differences similar to those in mice, which may have important implications for understanding sex-specific vulnerabilities to various human diseases.

## Discussion

Our findings highlight sex as an important physiological determinant of LVChP function, regulating its transcriptome, proteome, and secretome. Our molecular profiling and functional experiments highlight important sex differences in the LVChP, especially related to immune regulation and secretory signalling. Moreover sex differences are conserved in humans, and may underlie disease pathways.

Beyond its classical roles as a gatekeeping barrier and in CSF production, the LVChP is a dynamic source of signals that can influence multiple regions of the CNS. One of our key findings is the difference in LVChP secretory signalling between sexes. Proteomics revealed that both male and female LVChP secretomes contain diverse bioactive signalling molecules, including growth-promoting, developmental factors, hormones, neuropeptides, immunomodulatory mediators, enzymes and extracellular matrix proteins. While epithelial cells are a primary secretory cell type in the ChP, our analysis suggests that the LVChP secretory output could be shaped by contributions from several cell populations, and not just the epithelium. Interestingly, cell compartment analysis of male and female DEGs revealed distinct terms associated with secretory activity, including multiple vesicle related pathways and synaptic and neurotransmitter release related components in male and in estrous phase. As such, it will be key to explore how sex influences the distinct secretory modalities in the LVChP. Strikingly, secretory signalling differences between males and females have functional consequences, as highlighted by the sex-dependent effects on V-SVZ neural stem cells in vitro. ChP factors not only affect multiple cell types in the V-SVZ (Silva-Vargas et al 2016), but they can also modulate other brain areas via their delivery in the CSF (Baruch et al 2014, Phillipps et al 2025, Spatazza et al 2013). Increased CSF production itself has been reported in female mice (Liu et al 2020), however the mechanism is still not fully elucidated, as analysis of components of CSF production and transport machinery components in epithelial cells suggested overall similarity between sexes (Andreassen et al 2022). Interestingly, some of these components change with estrous cycle (Israelsen et al 2023). Further analysis will shed light on the complex regulation of CSF production and its dynamics, and how they can affect the brain environment more broadly.

The ChP is an immunological niche at the blood–CSF interface, hosting immune cells such as T cells and BAMs (Baruch et al 2013, Cui et al 2021) and sensing systemic and CSF-derived cues, to respond with tailored immune activation to acute and recurrent inflammation (Baruch et al 2014, Marques & Sousa 2015). Our transcriptomic and proteomic analyses suggest that sex is a major regulator of the LVChP immune landscape in mouse and humans. In females, upregulated pathways include cytokine and chemokine signalling, along with immunomodulatory factors, MHC expression, antigen presentation and adaptive immunity regulators in the secretome. These features could confer greater responsiveness to peripheral immune challenges but also a higher propensity to inflammatory response. In males, complement pathway components, central to pathogen recognition and inflammatory priming by the innate immune system (Schartz & Tenner 2020), are upregulated in the proteome in both mouse and human. These findings suggest a finely tuned, sex-specific balance between complement activators and inhibitors, shaping ChP immune surveillance and inflammatory responses across species that may influence inflammatory and neurodegenerative disease processes. Defining the balance of immune–secretory crosstalk between short-term plasticity and potentially harmful long-term consequences in both in health and disease will be crucial.

Intriguingly, our analysis also revealed striking sex differences in BAMs across compartments, with females having more stromal macrophages, and males more epiplexus cells. To our knowledge, this is the first report to indicate significant sex-related differences in LVChP cell types. Different morphologies of BAMs were more abundant in females, and not all epiplexus cells express P2YR12, suggesting further heterogeneity in BAMs. A key open question is the functional significance of BAM heterogeneity, and how this contributes to disease states. Indeed their compartment specific distribution and transcriptional heterogeneity likely shape macrophage interactions with distinct ChP cell types. Given the role of BAMS in pathogen defense and inflammation (Cui et al 2021), BAM subpopulations may be key modulators of ChP immune surveillance and broader CNS immune responses, with potential relevance to neuroinflammatory and neurodegenerative disease processes.

At the sensing level, the LVChP epithelial and stromal compartments express a broad repertoire of receptors and ion channels reflecting the role of the LVChP as a sensor of and responder to whole body states that dynamically adjusts its transcriptome and secretome to fine-tune its output in response to physiological changes. Intriguingly, in both mouse and human, female upregulated LVChP signatures suggest an enhanced ‘sensor’ capacity, with many pathways related to response to stimuli, and increased transcription factor activity induced by diverse stimuli including hormonal, oxygen, immune status, and metabolic cues, potentially reflecting a more dynamic adaptation to physiological shifts and increased sensitivity to signal dysregulation. LVChP signatures upregulated in males in both mouse and human were related to cilia, mechanotransduction and ion channels related terms. In the future it will be important to examine dynamic changes in receptor and ion channel repertoire expression and membrane associated structures that may modulate the sensing capacity of the LVChP.

The choroid plexus expresses multiple sex hormone receptors (Santos, et al. 2017), suggesting that several aspects of LVChP biology might be influenced by sex hormones. Interestingly, the LVChP shows global differences between estrous and diestrous at the transcriptomic level which may result in changes in CSF composition. Importantly, we show that some sex hormone receptors are enriched in the ChP as compared to brain areas, and that within the ChP itself, sex hormone receptor expression is not uniform across ChP cell types. For example, LVChP BAMs express several sex hormone receptors, suggesting that hormones fluctuations could impact their cytokine and chemokine profiles at different phases of the estrous cycle. This is consistent with change of chemokines and cytokine profiles observed upon gonadal removal in both sexes (Quintela et al 2013, Santos et al 2017). Importantly, sex hormones are also active in the male LVChP. For example, the Prolactin receptor (Prlr), which is upregulated in both female LVChP transcriptome and proteome, exhibits a dynamic expression pattern across estrous cycle and in various reproductive states in females, but is also active in males (Bridges et al 2011, Kalyani et al 2016, Pi & Voogt 2002). Future work will be essential to dissect how sex hormones act on distinct LVChP cell types in both males and females, and to pinpoint how sex chromosome related differences and hormonal cycles intersect to shape ChP biology.

In humans, changes in CSF secretion rates and composition are often altered in disease, with ChP derived proteins emerging as early disease biomarkers (Bothwell et al 2019). Immune-related proteins in the CSF derived from the ChP have been implicated in neurodegenerative and inflammatory diseases (reviewed by (Courtney et al 2025). These changes can be due to disruption of ChP cell–cell communication, whether through altered signalling, receptor expression, or secretome composition, of multiple ChP cell types. Defects in the secretory function of the ChP epithelium have also been described in multiple neurological disorders and are being re-examined as a potential drivers during the early steps of disease development (Nutter et al 2023, Rodriguez-Lorenzo et al 2020, Tanabe et al 2025, Yan et al 2025). A key next step is to resolve heterogeneity amongst different ChP cell types, and how define how these contribute to disease. One of the most striking sex differences is the strong upregulation in mouse and human LVChP and CSF of α2-macroglobulin (A2M), which has several important roles including protease inhibition and amyloid clearance, and may contribute to neurodegenerative pathogenesis, including Alzheimer’s disease (Qiu et al 2022).

The LVChP is an important sensory and secretory hub, integrating diverse signals from both peripheral and CNS compartments. Here we show that sex may be an important physiological determinant of immune, secretory, and homeostatic functions. Given the sex-biases of several neurodegenerative diseases, it will be important to understand whether the sex differences we have uncovered contribute to their prevalence. In the future, it will be exciting to dissect how sex hormonal regulation, immune, and metabolic inputs are orchestrated at the molecular level by each cell in the LVChP to shape the brain environment in physiology and contribute to resilience and vulnerability in disease.

## Methods

### Animal use

Two- to four-month-old male and female CD1 mice (Charles River Labs, Strain #:03257), *h*GFAP::GFP mice (Jackson Laboratory) (Zhuo et al 1997) and heterozygous CX3CR1-GFP mice (Jackson Laboratory, Strain# 005582)(Jung et al 2000) were used for our experiments. All mice were housed in a 12:12h light/dark cycle with *ad libitum* access to food. Estrous cycle status of females was assessed via vaginal swabs as described (Ajayi & Akhigbe 2020). All procedures were done under a license approved by the veterinary office of canton Basel-Stadt.

### Bulk RNA analysis of mouse and human transcriptome

For mouse bulkseq data, individual LVChPs from males and females in diestrus and estrus phases of the estrous cycle were isolated for transcriptome analysis (n=3 mice per group). RNA was extracted from individual LVChPs using the RNeasy Plus Micro Kit (Qiagen, Cat#74034) according to the manufacturer’s instructions. RNA sequencing was done by paired end sequencing with Illumina NovaSeq 6000 V3.4.4 at the Genomics Facility Basel. For human bulkseq data (Rodriguez-Lorenzo et al 2020), sequencing data were obtained from GEO repository (GSE137619). Only LVChP from healthy controls were selected for analysis.

The following human LVChP samples from the original study (as listed in GEO depository) were used for transcriptomic analysis:

GSM4083354 Control CP 1 – Female, age: 68
GSM4083355 Control CP 2 – Male, age: 55
GSM4083356 Control CP 3 – Female, age: 64
GSM4083357 Control CP 4 – Female, age: 57
GSM4083358 Control CP 5 – Female, age: 70
GSM4083359 Control CP 6 - Male, age: 59

The mouse and human sequencing files were then processed using nf-core pipeline “rnasplice” (https://nf-co.re/rnasplice/, version: 1.0.4). Briefly, the pipeline performs quality control with multiqc (v1.18) and trimming using TrimGalore! (v0.6.7). The human and mouse samples were mapped using STAR aligner (2.7.10a) to the human reference genome (Homo sapiens GRCh38 v.114) and the mouse reference genome (Mus musculus GRCm39.113), respectively and raw counts were determined using Salmon (v1.10.1). Raw counts were preprocessed by NoisyR (v 1.0.0) (Moutsopoulos et al 2021) to remove technical noise from sequencing data before being analyzed by DESeq2 (v1.49.3) (Love et al 2014) to determine differential expression and normalisation of the gene-wise count data. Genes with an adjusted p-value (using Benjamini-Hochberg method) lower than 0.05 were considered differentially expressed. For all downstream functional analysis DESeq2 significant DEGs were filtered to remove sex chromosome-associated genes (keeping only autosomal genes for analysis).

### Gene set enrichment analysis using GOAT algorithm of adult male and female LVChP transcriptome

Gene set enrichment analysis was performed using the GOAT algorithm (Koopmans 2024). The analysis was performed using the output of DESeq2 analysis of human or mouse transcriptome (filtered for autosomal genes only). The mouse DESeq2 output was then further modified to map mouse genes to human homologues followed by an addition of human Entrez IDs to the converted human gene symbols (to comply with the input file instructions as described on the GOAT website - https://ftwkoopmans.github.io/goat). Properly formatted input files were then imported for GOAT analysis using the available online tool (https://ftwkoopmans.github.io/goat). For both the human and mouse analysis, the same parameters were used:

**gene score type** – effect size
**gene set size minimum** – 10
**gene set size maximum** – 1500
**p-value adjustment –** Bonferroni (adj. p value – threshold 0.05)

The exception was the processing of the human transcriptome for the generation of the gene set similarity heatmap, where the gene set size minimum was set to 100 in order to obtain a reduced number of significant hits compatible with the heatmap generation requirements. The online GOAT tool was used to generate all GOAT analysis-related plots and GOAT analysis correlative heatmaps shown in the paper. Also the processed output was used in several pathway enrichment plots (Fig.1, Supplementary Fig.2, Supplementary Fig.3 and Fig.5, Supplementary Fig.6 respectively).

### Spatial transcriptomic analysis

Spatial analysis was performed using the publicly available MERFISH dataset from adult male and female brains (Zhang et al 2023) obtained from the CellxGene database (https://cellxgene.cziscience.com). The following raw and imputed datasets as listed in the database were used for downstream analysis:

**MERFISH raw data:**
WB_MERFISH_animal1_coronal (Female)
WB_MERFISH_animal2_coronal (Male)

**MERFISH imputed data:**
WB_imputation_animal1_coronal_anterior (Female)
WB_imputation_animal1_coronal_posterior (Female)
WB_imputation_animal2_coronal (Male)

Imputation was performed by Allen brain map consortium as described here: (https://github.com/ZhuangLab/whole_mouse_brain_MERFISH_atlas_scripts_2023/tree/main/scripts/integrate_MERFISH_with_scRNA-seq).

We used plotly (5.19.0) for visualization and manual selection of LVChP from the individual coronal sections of male and female brain included in the dataset. Next, manually annotated cells were extracted and pooled to obtain filtered dataset containing only LVChP cells. Cell type annotation information used in the paper was provided by Allen Brain Map institute using their reference single cell brain dataset (Yao et al 2023) and already included in the dataset. For more information on cell annotation see https://portal.brain-map.org/atlases-and-data/bkp/mapmycells. For spatial data visualization, either the original raw MERFISH data (referred to as “raw”) or the imputed MERFISH data (referred to as “imputed”) were used, as indicated in the manuscript. Prior to analysis and plotting, raw data were log2-normalized. For plotting cell type–specific gene expression, globally scaled imputed log2-transformed expression values were used, representing the mean gene expression in a given cell type relative to the mean expression across all analyzed cells. Dot plots were generated using Scanpy (v1.11.1).

#### Gene set enrichment analysis (GSEA) of adult male and female LVChP proteome

Proteome data from male and female LVChP was used to perform modified Gene Set Enrichment Analysis using fGSEA (Korotkevich et al 2021, Subramanian et al 2005). Analysis was performed using the Gene Ontology Biological process (GO:BP) database obtained from msdigbr R (https://igordot.github.io/msigdbr/) and using the KEGG database obtained from KEGGREST R (https://bioconductor.org/packages/KEGGREST). The following parameters were used:

**Statistical test performed:** fgseaSimple() with 1000 permutations
**stats**: A numeric vector named ranked_proteins containing protein t-statistic values, which serves as the pre-ranked list.
**minSize**: **15** (pathways with fewer than 15 genes were excluded).
**maxSize**: **500** (pathways with more than 500 genes were excluded).
**nproc**: **1** (single-threaded processing was used for reproducibility, as set by options(mc.cores = 1) and then nproc = 1).

GSEA was performed on the entire proteome data to identify significant differences in biological processes between the sexes. Top pathway terms were selected based on adjusted p value (using Benjamini-Hochberg method) and ordered for plotting based on NES value.

### ENRICHR analysis of mouse and human DEGs and DEPs

The ENRICHR online tool (https://maayanlab.cloud/Enrichr) was used to investigate the pathway enrichments for mouse transcriptome DEGs, mouse proteome DEPs (LVChP proteome DEPs, DEPs filtered for putative secreted factors (Wang et al 2024) or DEPs overlapping with CSF proteome (Gorska et al 2024) and human proteome DEPs (Wesenhagen et al 2022) (Chen et al 2013, Kuleshov et al 2016). The hits for plotting were selected based on p-value (determined using Fisher’s exact test) and plotted either using combined score or – log10(p-value) on the x-axis.

#### Transcription factor activity inference from bulkseq mouse and human transcriptomes

We used Decoupler R (Badia et al 2022) to infer the differential activation of transcription factors between female and male mouse and human transcriptomes. The analysis was performed following the instructions from the official online tutorial for bulkseq data processing (https://saezlab.github.io/decoupleR/articles/tf_bk.html). Briefly, the DESeq2 output (only with autosomal genes) from the analysis of mouse and human transcriptomes was used as input. CollecTRI database was used as reference for curated transcription factor (TF) regulatory network information. TF activity scores were determined using the Univariate linear model (ULM) method. For the TF inference, the stat value (corresponding to t-value referred to in the tutorial) from DESeq2 output was used. Decoupler was also used for visualization of the differential target gene expression for selected TFs.

### Proteomics

#### Sample preparation for Mass Spectrometry based proteome analysis

Mouse tissue samples were dissolved in lysis buffer containing 5% SDS, 100 mM TEAB, and 10 mM TCEP and heated at 95 C for 10 min. Lysates were further sonicated using a PIXUL™ Multi-Sample Sonicator (Active Motif) using default settings for 20 min. Carbamidomethylation of cysteins was performed by addition of 20 mM Iodoacetamide at 25 C in the dark for 30 minutes. Reconstituted eluates were subjected to S-Trap based digestion and purification according to the manufacturers procedure (Protifi.com). Dried peptides were dissolved in 0.1% aqueous formic acid solution at a concentration of 0.1 mg/ml prior to injection into the mass spectrometer.

#### Mass Spectrometry Analysis

Sample injection order was block randomized based on the experimental conditions. For each sample, a total of 0.15 μg peptide mixture was subjected to LC-MS analysis using an Orbitrap Eclipse Mass Spectrometer (Thermo Fisher Scientific) equipped with a nanoelectrospray ion source. Peptide separation was carried out using an UltiMate 3000 system (Dionex) equipped with a RP-HPLC column (75μm × 30cm) packed in-house with C18 resin (ReproSil-Pur C18-AQ, 1.9 μm resin; Dr. Maisch GmbH, Germany) and a custom-made column heater (60° C). The following gradient was used for peptide separation: from 2% B to 12% B over 5 min, to 30% B over 70 min, to 50 % B over 15 min, to 95% B over 2 min, followed by 18 min at 95% B. Buffer A was 0.1% formic acid in water and buffer B was 80% acetonitrile, 0.1% formic acid in water.

The Orbitrap Eclipse mass spectrometer was operated in data independent acquisition mode (DIA) with a total cycle time of approximately 2 seconds. Each MS1 scan was followed by high-collision-dissociation (HCD) of 42 DIA windows ranging from 400 m/z to 900 m/z using a fixed HCD normalized collision energy of 33%. AGC target for MS1 was set to 250% with a maximum accumulation time of 45 ms and a resolution of 120,000 FWHM at 200 m/z. For MS2, AGC target was set to 800% with a maximum accumulation time of 22 ms and a resolution of 15,000 FWHM at 200 m/z.

### Protein Identification and Quantification

DIA data analysis was performed using DIA-NN (v1.8.1) (Demichev et al 2020) with default settings. For database search, a target-decoy based strategy using the UniProt murine database (version 2022-Feb-02, including commonly known protein contaminants) was used. For DIA-NN settings, double-pass mode was selected for the neural network classifier protein names were selected for protein interference detection. Initial protein Quantification and statistical analysis was performed by the application of the MSstats (v4.8.6) (Choi et al 2014) based on the DIA-NN output. Limma (v3.62.2) linear models (lmFit) were applied to the log₂-transformed protein intensities from the MSstats output, with a +1 × 10⁻⁴ offset added to stabilize variance estimates. Contrasts were fitted using contrast.fit, and statistical inference was performed with robust empirical Bayes moderation (eBayes, robust = TRUE).

### Proteomic analysis of putative global or CSF-specific LVChP secretome

For preparing a reference list of secreted proteins for secretome analysis, we used list of Mouse secreted proteins from SEPDB database (Wang et al 2024) refered to as “Mouse Secreted Protein Datasets” (see https://sysomics.com/Download). To extract and generate list of secreted proteins for the comparative analysis with LVChP proteome a gene had to be listed either as “secreted” in the column UniprotEvidence or labelled as “TRUE” in the column “CM-Brain” in the input list of Mouse secreted proteins. This filtered list was than used to identify overlapping presumably secreted proteins in male and female mouse LVChP proteome. For identification of proteins from mouse LVChP proteome potentially secreted into CSF, we first prepared a list of proteins detected in mouse CSF (Gorska et al 2024). Any protein that was successfully detected in the CSF of all 3 wild-type animals was used to prepare a list of mouse CSF proteins for downstream overlap analysis with our male and female mouse LVChP proteome.

### Antibody arrays of LVChP conditioned medium

Male, female estrus, female diestrus and female pooled (estrus and diestrus pooled) LVChP conditioned medium samples were prepared as described in the *in vitro* culture LVChP Conditioned Medium section below. Proteome profiler mouse XL cytokine arrays (R&D systems, Cat#ARY028) were used according to the manufacturer’s instructions. Relative levels of the different proteins were calculated based on densitometry. These levels were log2transformed and the Wilcoxon rank-sum test was used to determine the p-value per protein (two levels of strictness applied p < 0.05 or p < 0.1). Benjamin-Hochberg test was applied to scale point sizes for slope-test plotting. Each antibody array was performed in triplicate using independent male, female mixed, female estrous and female diestrous samples.

### V-SVZ Single cell dataset processing

A previously published single cell dataset of male and female lateral ventricular-subventricular zone (V-SVZ) (Cebrian-Silla et al 2021) was used to obtain receptor expression profiles across V-SVZ cell types. Briefly, we obtained the raw counts and metadata from the GEO depository. These were used to generate a single cell object that contained individual cell type annotations. Ambient RNA was removed using SoupX (Young & Behjati 2020), doublets were identified using scDblFinder and Vaeda (Germain et al 2021, Schriever & Kostka 2023) and cells that were identified as doublets based on both methods were removed. This dataset was then used for downstream analysis.

### Ligand-receptor analysis

For ligand-receptor mapping, all proteins detected by antibody arrays and those overlapping between the LVChP sex-pooled proteome and the SEPDB database (Wang et al 2024) of secreted proteins or the CSF proteome (Gorska et al 2024) were combined to generate the input ligand list, to detect their corresponding receptors. Next, we used Liana (Dimitrov et al 2022) to map ligands to receptors using the mousecensus reference database. Using original cell type annotations (Cebrian-Silla et al 2021), we determined the expression levels of mapped receptors in selected cell subtypes (B-NSCs, A-NBs, C-TACs, Ependymal cells, Microglia, OPC-Oligo, where OPC-Oligo were originally labelled as OPC/Oligo). For each receptor, a z-score was calculated across all analyzed cell types. We used the following parameters to filter receptors for plotting:

Minimal number of cell types expressing the receptor: 2
Minimal fraction (percentage) of cells expressing given receptor : 1
Minimal Z-score threshold: 0.25

For the L-R dot plot, the hits are ranked based on Z-score variation plotted in descending order. Z-scores are normalized per receptor not globally.

### Human CSF analysis

For downstream analysis of sex-specific differences in human (Wesenhagen et al 2022) we used the list of proteins that were significantly different based on sex and that included the METSIM dataset in supplementary table S7 (Wesenhagen et al 2022)). We used the β value as fold changes of proteins between male and female CSF. The Shiny app available as part of this study (https://kwesenhagen.shinyapps.io/Aging_proteomics) was used to generate protein enrichment box plots in Supplementary Fig. 6j, k.

### Generation of semantic heatmap of related terms for human CSF analysis

The semantic heatmap shown in Fig 5g was generated as follows. The significantly enriched human CSF proteins were selected by using |logFC| 0.25 and padj > 0.05. These were used in EnrichR analysis using EnrichR (Kuleshov et al 2016) using the following databases:

- GO Biological Process 2025
- KEGG 2021 Human
- Reactome Pathways 2024
- Elsevier Pathway Collection

The top 20 hits based on non-adjusted p-value were selected from each database, pooled and sorted into semantic categories based on presence of the keywords listed below:

**Immune / Defense category:** immune, macrophage, antigen, defense, bacteria
**Coagulation category:** coagulation, hemostasis
**Complement category :** complement

For visualization purposes, the significance value (−log10(Adjusted P-value)) of the heatmap is capped at a maximum value of 25.

### qPCR analysis

RNA was extracted from LVChP using the RNeasy Plus Micro Kit (Qiagen, Cat#74034) according to the manufacturer’s protocol. cDNA was generated with the Promega kit (Cat#3802). Quantitative polymerase chain reaction (qPCR) was performed using SYBR green (Roche) for detection on a Light Cycler Instrument II (Roche) according to the manufacturer’s instructions in triplicate. Rpl13a was used as a housekeeping gene. The thermocycling programme consisted of 95°C for 10 minutes, followed by 40 cycles at 95°C for 15 seconds and 60°C for 1 minute.

Sequences of primers are as follows:

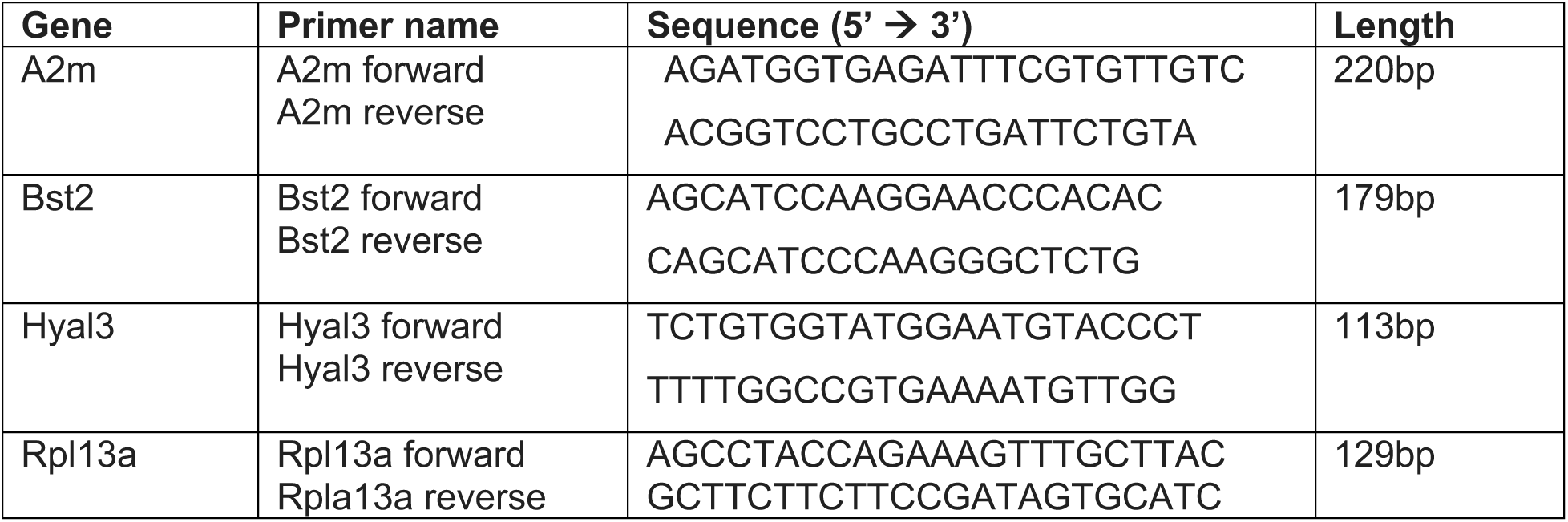

### Tissue preparation

Mice were deeply anaesthetised by intraperitoneal injection of Esconarkon and perfused with 0.9% saline followed by ice-cold 4% paraformaldehyde (PFA, Electron Microscopy Sciences) in 0.1M phosphate buffer. Brains were postfixed in 4% PFA for 24 hours at 4°C, and washed with 1X phosphate-buffered saline (PBS), transferred into a 15% sucrose solution followed by 30% sucrose solution overnight. Brains were embedded in OCT and stored at −80°C until use. 12 µm thick coronal sections were cut using a Leica CM30505 cryostat. For whole mounts of the LVChP, mice were perfused with 0.9% saline and lateral ventricle choroid plexuses dissected from the brain, fixed in ice-cold 4% PFA for 30 minutes and washed in 1X PBS.

### Immunostaining

#### Cryosections

Sections were thawed, washed in 1X PBS, and boiled for 15 minutes at 100°C in antigen retrieval solution (DAKO). Sections were washed and permeabilised in PBST (1X PBS supplemented with 0.5% Tween-20), followed by blocking in PBTA (1X PBS supplemented with 5% donkey serum, 0.3% Triton X-100 and 1% BSA), and incubated in primary antibodies in PBTA at 4°C overnight. After washing in PBST, sections were incubated with secondary antibodies in PBTA for 1 hour at room temperature, washed in PBST and counterstained with 4’, 6-diamidino-2-phenylindole (DAPI) (1:500). Slides were mounted with Aqua-Poly/Mount (Polysciences).

#### LVChP whole mounts

Whole mounts were permeabilised and blocked with 1X PBS containing 10% BSA and 0.2% TX-100 for 2 hours at room temperature, incubated in primary antibodies overnight at 4°C in 1X PBS containing 3% BSA and 0.2% TX-100, washed, and incubated in secondary antibodies for 2 hours at room temperatur, washed, counterstained with DAPI (Invitrogen, 1:500) and mounted on slides using Aqua-Poly/Mount.

#### Antibodies

The following antibodies were used for immunostainings: anti-AQP1 (mouse, 1:200, Santa Cruz, sc-55466), anti-AQP1 (rabbit, 1:100,Proteintech, 20333-1-AP), anti-NKCC1 (rabbit, 1:100, Cell Signaling, Cat#85403), anti-CSFR1 (sheep, 1:250, bio-techne, Cat#AF3818), anti-IBA1 (goat, 1:200, Abcam, Cat#ab5076), anti-IBA1 (rabbit, 1:250, IGZ Instruments AG (Fujifilm), Cat#019-19741), anti-GFP (chick, 1:250, Aves Labs, Cat#GFP-1020), anti-MRC1 (rabbit, 1:200, Abcam, Cat#ab64693), anti-P2RY12 (rabbit, 1:250, Alomone Labs, Cat#APR-012), anti-PECAM1/CD31 (rat, 1:200, BD Pharmington, Cat#550274). Secondary antibodies conjugated to Alexa Fluor 488, Alexa Fluor 568, Alexa Fluor 647 (Invitrogen) or to Cy3 (Jackson ImmunoResearch) were used at a dilution of 1:600.

#### RNAscope

RNAScope was performed according to the manufacturer’s protocol (ACD). Briefly, cryosections were washed in 1X PBS and baked for 30min at 60°C followed by fixation in 4% PFA for 15min at 4°C. After washing in PBS, sections were dehydrated in 50%, 70% and 2x in 100% EtOH for 5min each, incubated with peroxidase (RNAscope Multiplex Fluorescent Assay Kit (Advanced Cell Diagnostics)) for 10min at room temperature followed by antigen retrieval (Cat#322000) in a steamer at 100°C for 10min, washed with MilliQ dH2O, and permeabilized for 25min in Protease Plus (Cat. 322330) at 40°C. After washing in dH2O, slides were incubated with preheated (40°C) probes for 2 hours at 40°C. Sections were washed in wash buffer and stored in 5X SSC (Sigma-Aldrich, Cat#S6639-1L)) overnight at room temperature. Signal amplification (RNAscope Multiplex Fluorescent Detection Reagents v2, Cat#323110), was performed by incubating in RNAscope Amp-1 (30min, 40°C), AMP-2 (30min, 40°C) and RNAscope AMP-3 (15min, 40°C). Slides were then incubated in RNAscope Multiplex FL v2 HRP-C1 for 15min at 40°C, washed and the TSA 520 fluorophore (7523/1 KIT) diluted in TSA buffer (1:5000, 322810) against the C1 probe was incubated for 30min at 40°C, washed and then incubated with RNAscope Multiplex FL v2 HRP blocker for 15min, 40°C. After 2 washes in wash buffer, either the RNAscope Multiplex FL v2 HRP-C2 or RNAscope Multiplex FL v2 HRP-C3 (depending on the probe) were added to the slide and incubated for 15 min at 40°C, followed by washes and incubation with the TSA 650 fluorophore (7527/1 KIT) diluted in TSA buffer (1:5000) for 30min at 40°C. After washing, RNAscope Multiplex FL v2 HRP blocker was added to the slide and incubated for 15min at 40°C followed by washes. Finally, sections were stained for DAPI for 30s and mounted in Aqua-Poly/Mount (Polysciences). The following RNAScope probes were used: A2m (C1, acdbio, 853411), Bst2 (C2, acdbio, 502401-C2), Bscl2 (C2, acdbio, 1222851-C2), Cpxm2 (C1, acdbio, 559751), Itih5 (C1, acdbio, 460971), Gh (C1, acdbio, 44536).

#### RNAscope combined with immunofluorescence

For RNAscope with immunostaining we followed manufacturer’s instructions as described in RNAscope Multiplex Fluorescent Reagent Kit v2 User Manual (https://acdbio.com/documents/product-documents). Briefly, slides were processed as described for standard RNAscope until the antigen retrieval step that is performed with the co-detection target retrieval reagent (Cat#323165). Next, following the washing steps with fresh PBST, we performed overnight incubation of slides at 4°C with primary antibodies diluted in Co-Detection Antibody Diluent (Cat#323160). The next day, following a washing step with PBST, we followed the standard RNAscope protocol. After RNAscope hybridization, we incubated slides with secondary fluorescent antibody diluted in Co-Detection Antibody Diluent (Cat#323160). Finally, sections were stained for DAPI for 30s and mounted in Aqua-Poly/Mount (Polysciences).

### *In vitro* experiments

#### Control minimal medium

Control minimal medium was prepared as described in Silva-Vargas et al.2016 (Silva-Vargas et al 2016). It consists of DMEM/F12 medium supplemented with 0.6% glucose (Sigma), 0.1% NaHCO3 (Sigma), 5mM Hepes (Sigma), 2mM L-Glutamine (Gibco BRL), 50U/mL penicillin/ streptomycin (Gibco) and 1X hormone mix (10x) [DMEM/F12 supplemented with 0.5% glucose, 0.09% NaHCO3, 4mM Hepes, 0.8mg/mL apo-t-transferrin (Sigma), 0.02mg/mL insulin (bovine, Sigma), 90µg/mL putrescine (Sigma), 160nM progesterone (Sigma) and 240nM Na2SeO3 (Sigma)]. When used, EGF was added at 20ng/mL EGF (human recombinant, GibcoBRL).

#### LVChP Conditioned Medium

To prepare LVChP conditioned medium, 5 dissected LVChP were cultured in a transwell (Corning, Cat#3470) containing minimal medium without addition of progesterone for 6 hours at 37°C. The transwell containing the LVChPs was then removed and the medium collected from the well plate. Next, progesterone, at a final concentration of 160nM, was added to the harvested LVChPsec and the LVCHPsec was used either directly or stored at −80°C for later use (Silva-Vargas et al 2016).

#### In vitro NSC cultures

Activated NSCs were FACS purified from the V-SVZ of 2-3 month old *h*GFAP::GFP (Jackson Laboratory) as described (Codega et al 2014). FACS purified activated NSCs were centrifuged at 1300 rpm for 10 min at 4°C and resuspended in minimal medium. Cell number and viability were determined using a haemocytometer and Vybrant Dye staining (1:1000, Invitrogen). Activated NSCs were plated at a density of 1 cell/µl in ultra-low attachment 96-well plates (Corning) in male or female LVChP conditioned medium, minimal medium, or minimal medium +EGF. Activated clones were counted two days after plating. For all culture experiments, cells and LVChP conditioned medium were prepared from pooled females at different phases of the estrous cycle.

#### Imaging and quantification

Images were acquired using the Olympus confocal spinning disk microscopes SpinSR (CSU-W1) and SpinD (CSU-W1) using a UPL S APO 30x/ 1.05NA objective with Silicon oil. Tile scans of the entire LVChP in cross-sections or whole mounts were acquired, with z-stacks encompassing the entire thickness. Individual z step sizes were 0.41-1µm, depending on the staining. Images of the same immunostainings were acquired on the same microscope with the same laser power and exposure. Individual tiles acquired on the SpinSR and SpinD were stitched using the Fiji plugin *recursive Olympus vsi stitcher*. Images of cultured NSCs were acquired using the Leica DMi8 system at 10x.

#### Analysis of cross-section immunostainings

Maximum intensity projections were used to quantify coronal cross sections. The number of IBA1+, MRC1+, and P2RY12+ cells was counted using the *cell counter* plugin in ImageJ. The total number of cells per LVChP cross-section was determined using DAPI staining in QuPath2.3. The total number of macrophages along the entire dorsal-ventral axis of the LVChP per section was quantified in five coronal sections per mouse spanning the rostro-caudal axis of the LVChP (n=4-8 mice per immunostaining).

#### Analysis of whole mount immunostainings

The number of IBA1^+^ cells was counted using the *cell counter* plugin in ImageJ. Volumetric measurement of LVChP tissue was performed using surface rendering in Imaris. The surfaces tool was used to reconstruct the LVChP in three dimensions. Surface boundaries were defined based on fluorescence intensity thresholding. The total tissue volume for each LVChP was calculated automatically by Imaris from the rendered surface model. The LVChP was divided into head (rostral), mid and tail (caudal) regions, and three to four tiles were quantified in each of these three regions in three female and four male samples, respectively. IBA1^+^ cells with different morphologies were manually counted using the *cell counter* plugin and LVChP tissue volume was measured in Imaris using surface rendering.

BAM quantification in the entire LVChP was performed in CX3CR1-GFP mice. The IJ rolling ball algorithm was used for background subtraction to reduce noise. A median filter was applied and Otsu thresholding used to create masks for macrophages and small objects were filtered out, and the volume of segmented macrophage regions measured. To normalize to tissue volume, a Gaussian filter was applied to a separate channel and segmentation was done using a manual threshold. Finally, the volume of the tissue was measured.

#### Statistical analysis

Statistical analyses and graphs were performed using GraphPad Prism software (version 7). Significance was established using two-tailed student’s t-tests for pair-wise comparisons. 2-way ANOVA was used to assess the effect of male and female LVChPsec cultured with sex-matched or sex-swapped NSCs, followed by Šídák’s multiple comparisons test (Figure 3.i). In Supplementary Fig. 2e, significance was calculated using the adjusted p-value <0.05. In Fig. 4g and Supplementary Fig. 5f two-tailed student’s t-tests with a p-value <0.05 were considered upregulated.Significance was established at *p<0.05, **p<0.01, ***p<0.001, and ****p<0.0001. Error bars indicate the standard error of the mean (SEM).

### Datasets

Human bulkseq transcriptomic data (Rodriguez-Lorenzo et al 2020) were obtained from GEO depository (GSE137619).

SVZ single cell data (Cebrian-Silla et al 2021) were obtained from GEO depository (GSE165554).

The human CSF data were obtained from supplementary table S7 (Wesenhagen et al 2022). The spatial transcriptomic MERFISH dataset (Yao et al 2023) was obtained from CellxGene public database (collection ref: 0cca8620-8dee-45d0-aef5-23f032a5cf09).

All original datasets will be deposited on GEO.

## Acknowledgements

We thank Stella Stefanova and Janine Bögli from the Biozentrum FACS core facility, Laurent Guerard from the IMCF, Thomas Bock from the proteomics core facility, and SciCORE at the University of Basel. We thank Elena Hui for comments on the manuscript. This work was supported by Swiss National Science Foundation 31003A_163088 (F.D.), Swiss National Science Foundation 310030_208009 (F.D.), European Research Council Advanced Grant (No 789328) (F.D.), and the University of Basel.

