## Supplementary Information for "Choroid plexus sex differences in secretory signalling and immune compartments"

**Supplementary Fig 1. Pathway enrichment analysis of male and female lateral ventricle choroid plexus**

**a** Lollipop plot showing top 30 statistically most significant pathway terms from Gene Ontology – Molecular Function database analysis for male and female LVChP transcriptome using the GOAT algorithm. **b** Gene set similarity heatmap with Gene Ontology – Molecular Function pathway terms grouped into categories. Color labels in legend refer to overarching group categories and their enrichment in males (blue) or females (red). Similarity index indicates pathway term similarity. **c** Lollipop plot showing top 30 statistically most significant pathway terms from Gene Ontology – Cellular Component database analysis for male and female LVChP transcriptome using the GOAT algorithm. **d** Gene set similarity heatmap with Gene Ontology – Cellular Component pathway terms grouped into categories. Color labels in legend refer to overarching group categories and their enrichment in males (blue) or females (red). Similarity index indicates pathway term similarity. **e** Volcano plot illustrating the differential expression of target genes for transcription factor Stat5a between female and male LV ChP. The x-axis represents the log-fold change (logFC), and the y-axis shows significance (−log10(p−value)). Genes are colored based on consistency between the gene's regulation direction with the expected transcription factor's mode of regulation: red for consistent, blue for inconsistent.

**Supplementary Fig 2. Validation of DEGs in the LVChP**

**a** PCA plot showing all male and female LVChP samples. **b** Volcano plot of all male and female LVChP DEGs, including sex- chromosome associated DEGs. Significance thresholds adjusted p-value (padj) < 0.05, |log2FC|>0.25. **c** Heatmap showing expression of all sex chromosome-associated DEGs (z-scores of mean expression). **d** Combined dot plot showing top 5 hits for enriched cell type pathway terms from PanglaoDB Augmented 2021 database. Red dots show female-enriched terms and blue dots male-enriched terms. **e** qPCR validation of differential expression of selected DEGs. **f** *In situ* hybridization (RNAscope) of male and female enriched DEGs in LV ChP. Insets show higher-magnification images. Scale bar: 250µm. Inset scale bar: 10µm. **g, h** Immunostaining combined with *in situ* hybridization (RNAscope) for *A2m* in the 3$^{rd}$ (g) and 4$^{th}$ (h) ventricle ChP. AQP1 is an apical ChP epithelial marker. Scale bar: 10µm.

**Supplementary Fig 3. Analysis of sex hormone mediated signaling in LVChP**

**a** Spatial UMAP plots showing imputed spatial expression for selected sex hormone receptors expressed in LVChP. **b** Spatial UMAP plot showing imputed spatial expression for *Esr1*. Inset highlights enrichment of *Esr1* in non-epithelial cell types. **c** Volcano plot of differentially expressed genes (DEGs) in the LVChP of estrus and diestrus females. Significance thresholds adjusted p-value (padj) < 0.05, a |log2FC|>0.25. The top 5 most significant genes for estrus (green) and diestrus (violet) are highlighted. **d, e, f** Lollipop plots showing top 30 statistically most significant pathway terms from Gene Ontology (Biological Process – BP (**d**), Cellular Component - CC (**e**) and Molecular Function – MF (**f**)) database analysis for estrus and diestrus LV ChP transcriptome using the GOAT algorithm. Estrus hits in green, diestrus hits in purple. **g** Bar plot of inferred differential activity for top 40 most differentially active transcription factors (TFs) between estrus and diestrus. The x-axis represents the activity score, a measure of differential TF activity. Green bars highlight TFs with higher predicted activity in estrus, and purple in diestrus.

**Supplementary Fig. 4 Characterization of LVChP secretome ligands and V-SVZ receptor binding partners**

**a** Venn diagram showing overlap between total LVChP proteome (male and female LVChP combined) and CSF proteome (from Gorska et al 2024). Pie chart shows proportions of overlapping putative CSF secreted proteins significantly enriched or not in male and female LVChP proteome. **b** Combined dot plots showing pathway enrichment analysis for sex-enriched protein hits using Bioplanet_2019 database. Red and blue gradients (–log10(p-value)) indicate enrichment in females and males, respectively. **c** Bar plot showing sex-enriched hits from antibody array analysis of female and male LVChP conditioned medium (hits with p-value of <0.05 and <0.1). **d** Slope plot showing general enrichment of LVChP secreted proteins detected via antibody arrays. Green lines highlight estrus enriched proteins, violet lines diestrus enriched proteins. Darker color shade indicates enriched proteins (p value < 0.1). **e** Bar plot showing hits from antibody array analysis of estrus and diestrus LVChP conditioned medium (hits with p-value <0.1). **f** Schematic representation of coronal brain section showing LVChP in the ventricle and adjacent ventricular-subventricular zone (V-SVZ). Magnified view of V-SVZ shows the main cell types including ependymal cells (E), quiescent neural stem cells (qNSCs), activated NSCs (aNSCs), transit amplifying cells (TACs), neuroblasts (NBs), blood vessels (BVs), microglia and oligodendrocytes (oligo) and oligodendrocyte progenitors (OPC). **g** Ligand-receptor analysis showing receptor expression in different V-SVZ cell types (as annotated in Cebrian-Silla et al 2021) that potentially bind ligands from LVChP (based on pooled ligands from LVChP Secretome, CSF and antibody array analysis). Threshold used for selection of shown receptors: Minimal number of receptor expressing cell types: 2, minimal fraction of expressing cells in given cell type: 1%, minimal expression Z-score threshold variations across at least two cell types: 0.25.

**Supplementary Fig. 5 Characterization of border-associated macrophages in the LVChP**

**a** Representative image of a cross-section of CX3CR1-GFP mouse immunostained for GFP (green, CX3CR1$^{+}$ cells), IBA1 (magenta) and CSF1R (white). Scale bar: 50µm. **b** Overlap of pan-macrophage markers in the LVChP. **c,** Representative images of male and female LVChP whole mounts from CX3CR1$^{GFP}$ mice and corresponding quantification of CX3CR1 positive cells. **d** Representative image of whole mount preparations immunostained for IBA1 (cyan) and CD31 (red) showing macrophages associated with blood vessels. Box shows region shown at higher-magnification. Scale bar: 10µm. **e** Quantification of male and female total IBA1$^{+}$ BAMs in cross-sections. **f** Quantification of proportions of MRC1$^{+}$ and MRC1$^{-}$ stromal BAMs in the LVChP. **g** Dot plot of imputed spatial cell type-specific expression of *Irf8* in LVChP. **h** Quantification of overall LVChP proportions of P2RY12$^{+}$ and P2RY12$^{-}$ IBA1+ epiplexus cells.

**Supplementary Fig. 6 Characterization of human LVChP and disease pathway analysis**

**a, b** Combined dot plot showing top hits for enriched disease pathway terms from JENSEN_Diseases_Curated_2025 database in mouse LVChP putative secretome (**a**; filtered using SEPDB database) or CSF secretome (**b**; filtered for published CSF proteome). Red dots represent enriched terms in mouse female LVChP secretome and blue dots represent enriched terms in male. **c** Gene set similarity heatmap with Gene Ontology – Biological process pathway terms grouped into categories. Color labels in legend refer to overarching group categories and their enrichment in males (blue) or females (red). **d, e** Lollipop plots showing top 40 statistically most significant pathway terms from GO:CC and GO:MF databases analysis for human male and female LVChP transcriptome using the GOAT algorithm. **f** Bar plot displaying the inferred differential activity for the top 40 most differentially activated transcription factors (TFs) between human female and male LVChP transcriptome. The x-axis represents the activity score, a measure of differential TF activity. Red bars highlight TFs with higher predicted activity in females, while blue bars show TFs with higher predicted activity in males. **g** Lollipop plot showing top 40 statistically most significant pathway terms from Jensen_disease_curated_2025 database for human male and female LVChP transcriptome using the GOAT algorithm. **h** Volcano plot of sex-enriched proteins in human CSF between females and males. Significance thresholds were an adjusted p-value (padj) $< 0.05$ and a $|\log2FC|>0.25$. The top 5 most significant genes are highlighted for females (red) and males (blue), with some additional manually added hits displayed (A2M). **i** Heatmap comparing selected proteins in mouse LVChP secretome and human CSF. Center columns show log2FC (Female vs Male; red = higher in females, blue = higher in males); side columns show significance as −log10(padj) for Mouse (left) and Human (right) and enrichment in given sex. Genes are ordered by mouse effect size. **j** Box plot depicting Z-score values of proteins enriched in females, measured in cerebrospinal fluid (CSF) samples from male and female participants of the human CSF study (Kwesenhagen *et al.*). These proteins are also significantly enriched in the putative female lateral ventricle choroid plexus (LVChP) secretome. **k** Box plot depicting Z-score values of proteins enriched in males, measured in cerebrospinal fluid (CSF) samples from male and female participants of the human CSF study (Kwesenhagen *et al.*). These proteins are also significantly enriched in the putative male lateral ventricle choroid plexus (LVChP) secretome.

### Supplementary Figure 1

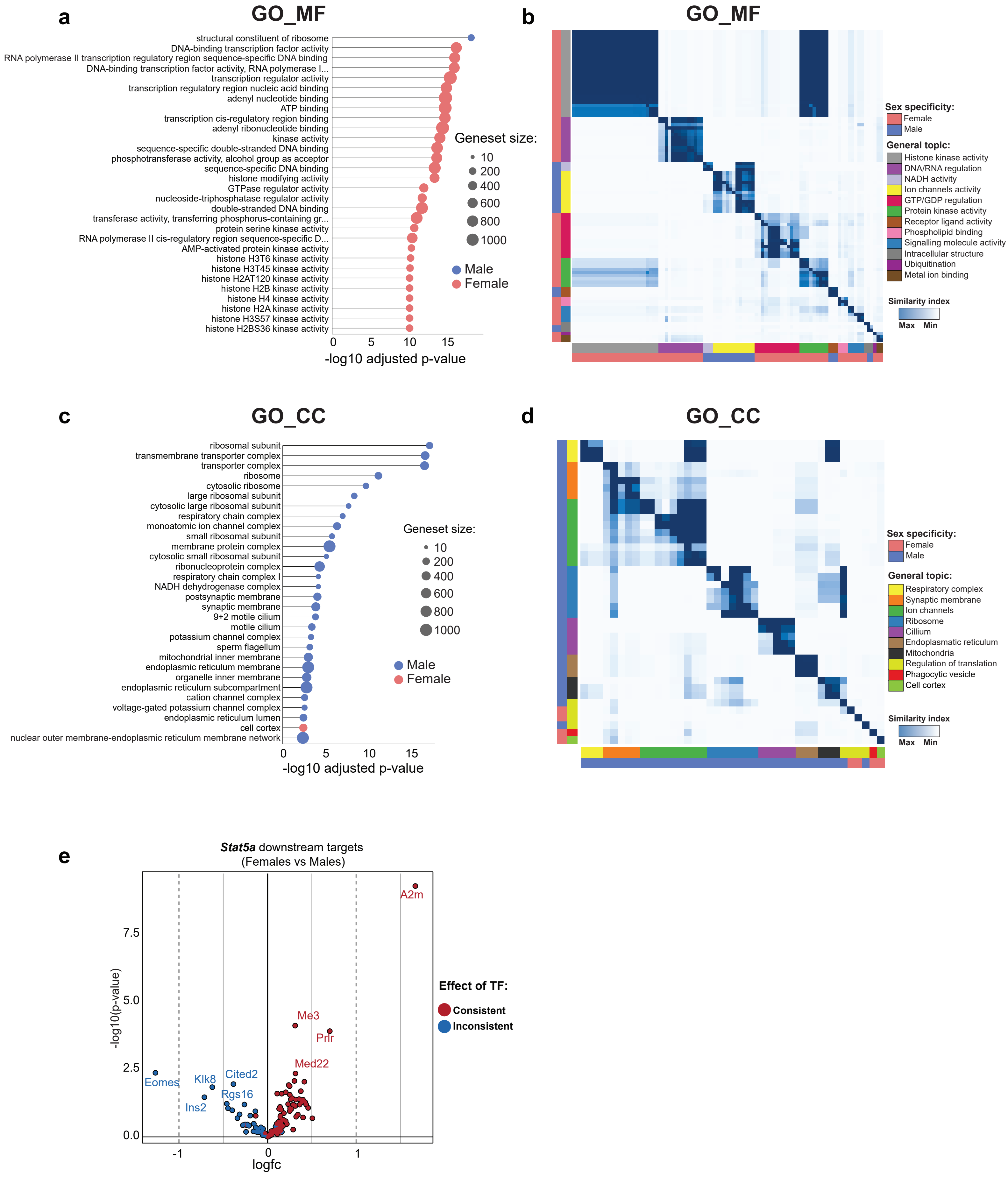

### Supplementary Figure 2

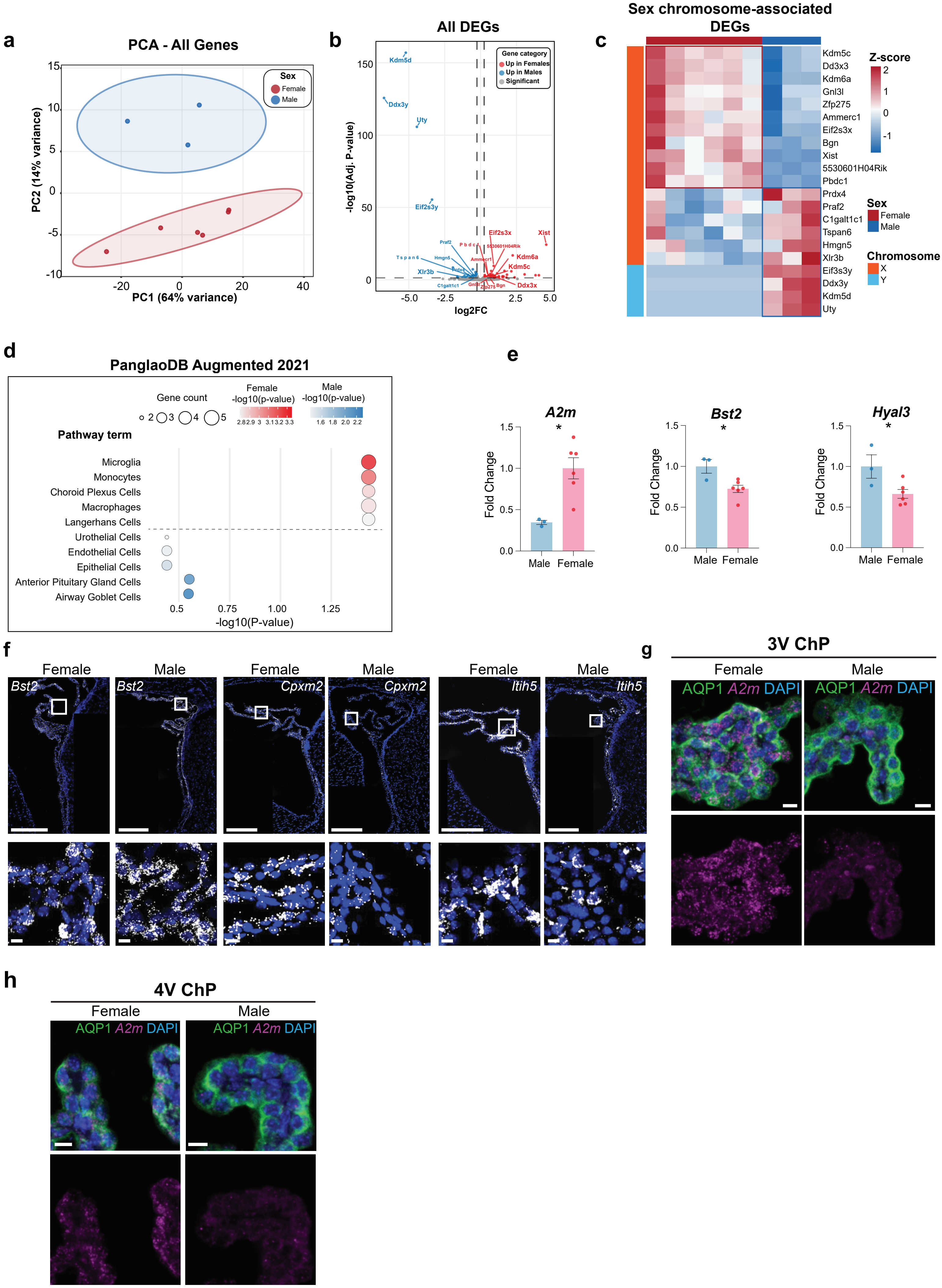

### Supplementary Figure 3

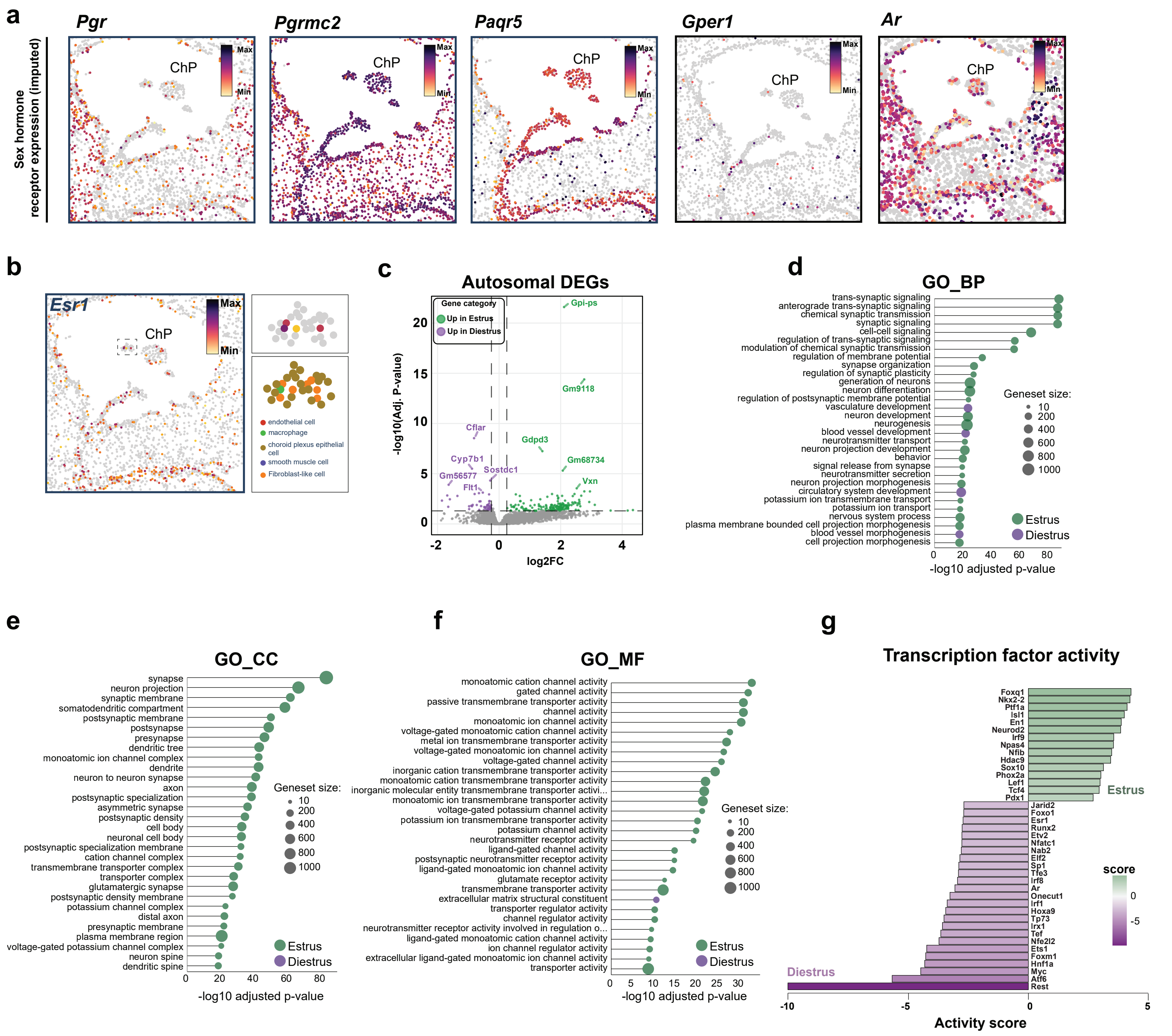

### Supplementary Figure 4

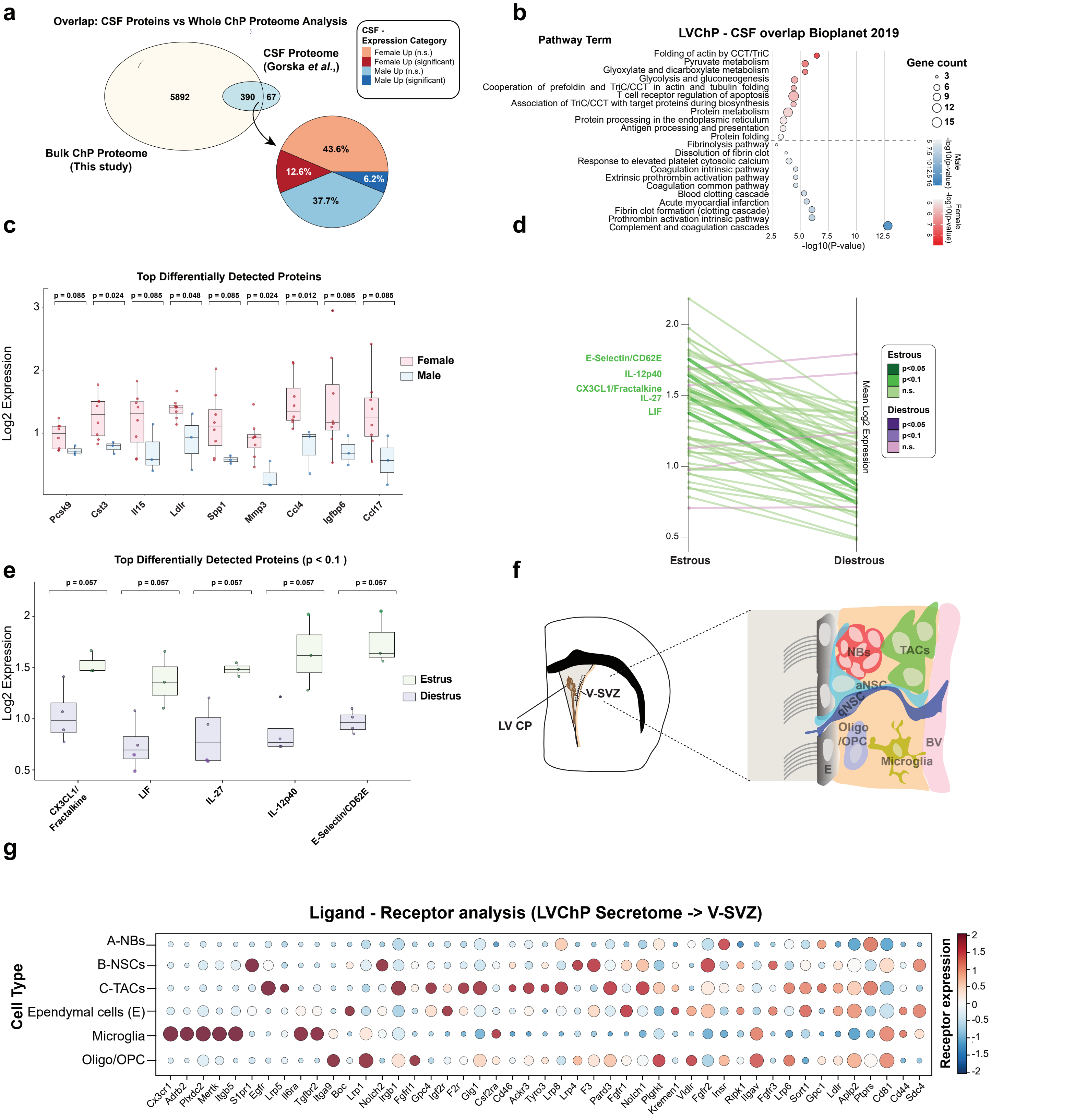

### Supplementary Figure 5

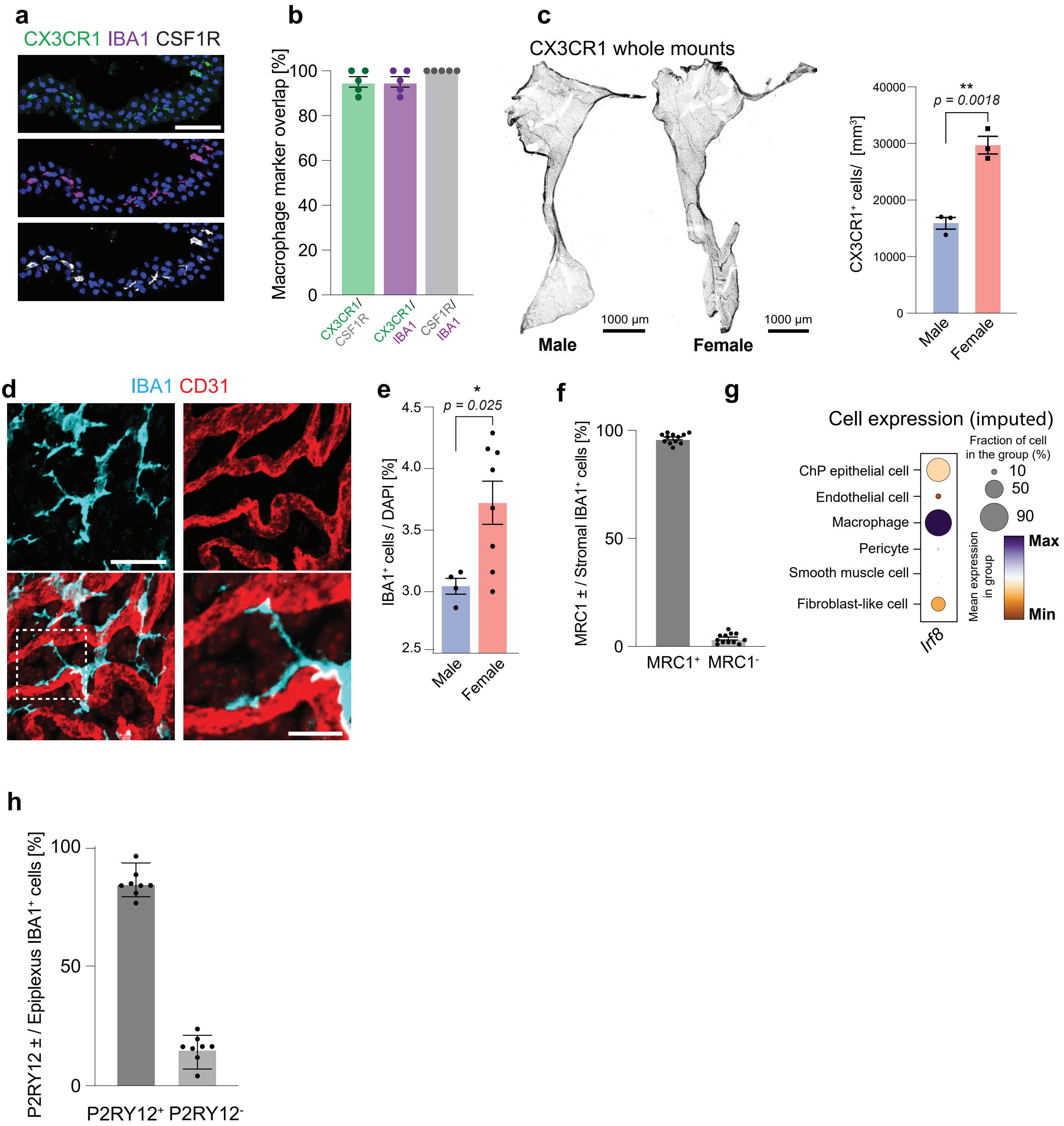

### Supplementary Figure 6

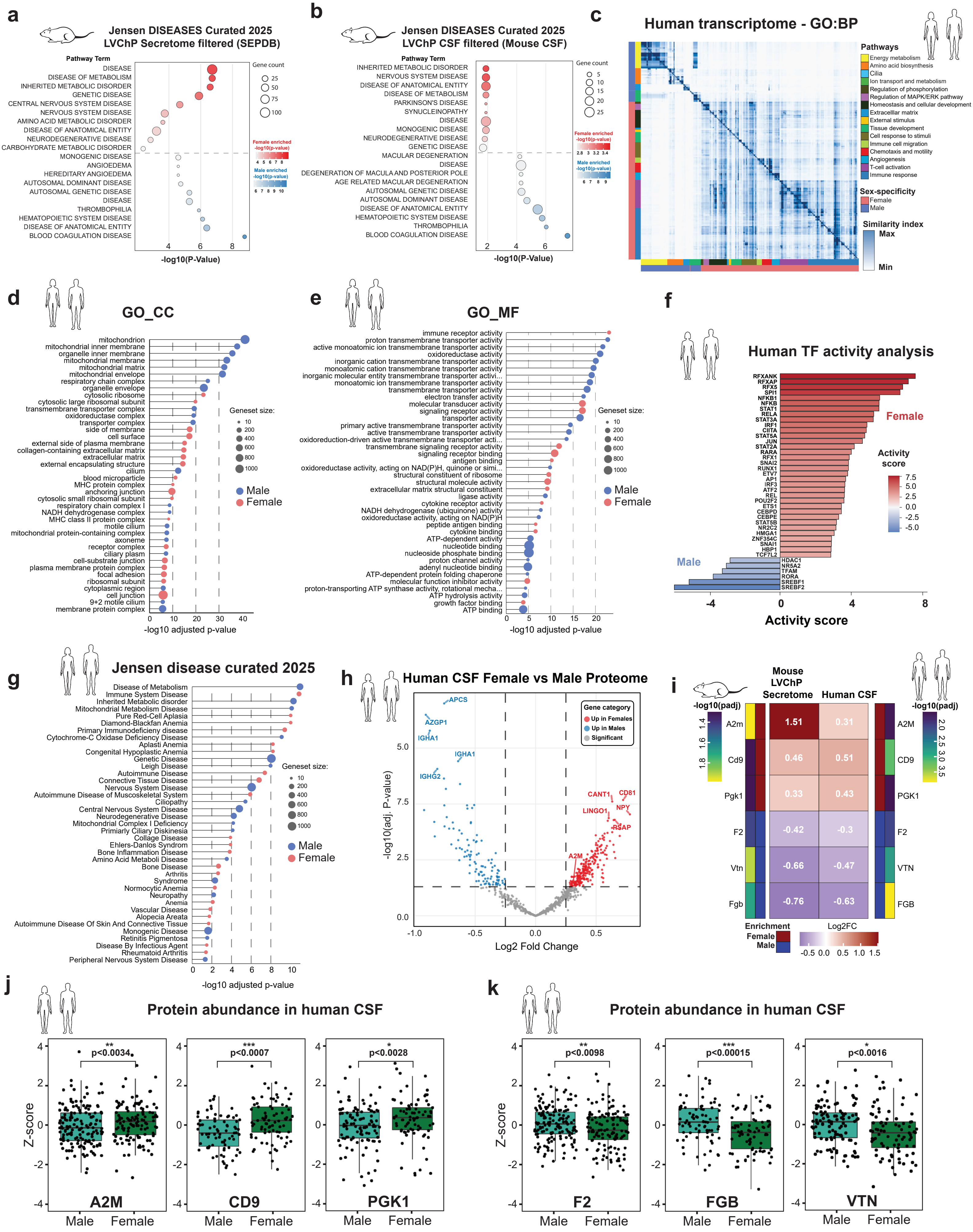
